# Transcriptome profiling of tendon fibroblasts at the onset of embryonic muscle contraction reveals novel force-responsive genes

**DOI:** 10.1101/2022.08.31.506026

**Authors:** Pavan K Nayak, Arul Subramanian, Thomas F Schilling

## Abstract

Mechanical forces play a critical role in tendon development and function, influencing cell behavior through mechanotransduction signaling pathways and subsequent extracellular matrix (ECM) remodeling. Here we investigate the molecular mechanisms by which tenocytes in developing zebrafish embryos respond to muscle contraction forces during the onset of swimming and cranial muscle activity. Using genome-wide bulk RNA sequencing of FAC-sorted tenocytes we identify novel tenocyte markers and genes involved in tendon mechanotransduction. Embryonic tendons show dramatic changes in expression of Matrix Remodeling Associated 5b (mxra5b), Matrilin 1 (matn1), and the transcription factor Kruppel-like factor 2a (klf2a), as muscles start to contract. Using embryos paralyzed either by loss of muscle contractility or neuromuscular stimulation we confirm that muscle contractile forces influence the spatial and temporal expression patterns of all three genes. Quantification of these gene expression changes across tenocytes at multiple tendon entheses and myotendinous junctions reveals that their responses depend on force intensity, duration and tissue stiffness. These force-dependent feedback mechanisms in tendons, particularly in the ECM, have important implications for improved treatments of tendon injuries and atrophy.

## Introduction

All cells experience mechanical forces from their environments, from adhesive interactions between adjacent epithelial cells to structural interactions with the surrounding extracellular matrix (ECM). A key question is how cells adapt and respond to force by modifying their local microenvironment. Force-responsive cellular mechanisms have been implicated in cell differentiation (D’Angelo et al., 2011), morphogenesis (Hamada, 2015; Keller et al., 2008), tissue maintenance and repair (Riley et al., 2022; Zhang et al., 2022). However, these mechanisms remain understudied in vivo, particularly those that involve cell-ECM interactions. Dramatic examples include tendons and ligaments of the musculoskeletal system. Tendons experience a broad range of contractile forces from muscles, such as extreme stretching forces on the human Achilles tendon during exercise, and their constitutive fibroblast populations (called tenocytes) constantly remodel the surrounding ECM to adapt (Subramanian & Schilling, 2015; J. H.-C. Wang, 2006). Tendon injuries and atrophy with aging are very common, and a better understanding of the roles of force in tendon development will aid in developing effective treatments.

Tendons are ECM-rich structures that connect muscles to cartilages and bones as well as to softer tissues. The events leading to the proper formation of their attachments relies largely upon cell-ECM interactions (Schweitzer et al., 2010; Subramanian & Schilling, 2015). For example, in the embryonic zebrafish trunk, myotendinous junctions (MTJs) at the vertical myosepta (VMS) of developing somites form via distinct tendon-independent and tendon- dependent stages (Subramanian & Schilling, 2015). In the tendon-independent phase, myofibers differentiate and secrete ECM proteins such as Thbs4b that localize to the pre-tendon ECM and mediate initial fiber attachment. This coincides with tendon progenitor cell (TPC) migration into the MTJ. Later, in response to muscle contraction, TPCs differentiate into mature tenocytes and extend long microtubule-rich processes laterally into the surrounding ECM of the VMS, with which they regulate ECM composition locally in response to force (McNeilly et al., 1996; Pingel et al., 2014; Subramanian et al., 2018). Contractile forces acting on these MTJs activate Transforming Growth Factor β (TGF-β) signaling in TPCs (Berthet et al., 2013; Pryce et al., 2009; Subramanian et al., 2018). Although not necessary for TPC specification, TGF-β induces expression of the transcription factors Scleraxis (Scx) and Mohawk (Mkx), which drive tenocyte fate by directly promoting transcription of collagens (i.e. Col1a1, Col1a2, Col12a1 and Col14) enriched in tendon ECM (Berthet et al., 2013; Maeda et al., 2011).

Cell type and ECM composition differ along the length of many tendons to aid in load bearing and force transmission. For example, the enthesis region where a tendon attaches to cartilage or bone is structurally graded in stiffness with fibrocartilage closer to the bone. This helps buffer mechanical stress between the elastic tendon tissue and rigid bony matrix (Lu & Thomopoulos, 2013). Fibrocartilage cells co-express Scx and Sox9, both direct transcriptional regulators of collagens, and muscle activity regulates the ratio of their expression levels (Blitz et al., 2013; Subramanian et al., 2023; Zelzer et al., 2014). This changes collagen levels, fibril size and organization during injury or repair, as has been shown both in vitro and ex vivo (Ireland et al., 2001; Pingel et al., 2014). We have also shown that muscle contraction is required for embryonic tenocyte maturation, morphogenesis and ECM production in zebrafish tendons in vivo (Subramanian et al., 2018; Subramanian & Schilling, 2014).

To identify genes regulated by muscle contraction in tendons we have performed genome-wide bulk RNA-sequencing (RNA-seq) on FAC-sorted tenocytes of zebrafish embryos during the onset of muscle contractions and active swimming behavior. In addition to upregulation of known tenocyte markers, we find several other genes up- or downregulated as tendons differentiate that have not been implicated in tenocyte development or mechano- transduction. These include genes encoding two ECM proteins, Matrix Remodeling Associated 5b (*mxra5b)* and Matrilin 1 (*matn1)*, as well as the transcription factor Kruppel-like factor 2a *(klf2a).* We confirm that muscle contraction regulates their transcription in tenocytes at later stages, after the onset of cranial muscle activity, by comparing wild-type and paralyzed embryos. Using genetic and physiological perturbations of muscle contraction in vivo, we show gene expression changes both in whole embryos and sorted tenocytes. Quantitative in situ methods show that their expression is contained within embryonic tendon entheses and MTJs and that their transcriptional responses to force vary depending on the strength and continuity of muscle contraction. These findings provide insights into tendon attachment specific and force-dependent feedback mechanisms in tendons during development in vivo, which have important implications for improved treatments for tendon disease, injury and atrophy.

## Results

### Onset of active muscle contraction alters tenocyte gene expression

We previously showed that trunk tenocytes in zebrafish undergo dramatic morphological transformations when muscle contractions begin (Subramanian et al., 2018, 2023). These occur when embryos transition from twitching (36 hours post-fertilization, hpf) to free-swimming behaviors (48 hpf), as well as between sporadic jaw contractions at 60 hpf, and free-feeding behavior at 72 hpf **(Fig, 1A-D)**. These morphological changes likely reflect force-induced transcriptional changes in tenocytes, in addition to changes driving differentiation. To identify potential force-responsive factors, we conducted RNA-seq with FAC-sorted populations of *Tg*(*scxa:mCherry)*-positive tenocytes isolated from dissociated twitching (36 hpf) or free-swimming embryos (48 hpf). The *Tg*(*scxa:mCherry)* line predominantly labels both embryonic trunk and cranial tenocytes. We FACS-sorted mCherry+ cells using WT stage-matched non-fluorescent embryos as negative controls **(Fig. 1—figure supplement 1).** Differential expression analysis revealed 2,788 differentially expressed genes (DEGs) between twitching and free-swimming stages with p-value < 0.05 **(Fig. 1E**) **(Supplementary File 1).** These included known tenocyte markers such as *tnmd, mkxa,* and *egr1* upregulated at swimming **(Fig. 1E-F)**, confirming that many of the sorted mCherry+ cells were tenocytes or TPCs. Principle Components associated with biological replicates segregated according to experimental condition (36 versus 48 hpf), validating library preparation **(Fig. 1G**) **(Fig.1—figure supplement 1)**. GO analysis for Biological Process (BP) terms associated with the top DEGs showed significant enrichment for “skeletal system development” and “ECM organization” **(Fig 1H)**. Surprisingly, these included *col2a1a* and *col9a1a*, which are typically associated with cartilage development and morphogenesis **(Fig. 1F)** suggesting that an early subset of *scxa+* cells in embryonic tendons are specified as developing enthesis cells (Subramanian et al., 2023). Dual-expressing *scxa/sox9a+* cells localize to cartilage attachment sites of cranial muscles at 48 hpf, prior to the onset of jaw movements **(Fig. 1A-D)**, consistent with specification of enthesis progenitors before the tendons or their skeletal muscle attachments become functional. These results also fit with recent single-cell sequencing studies of enthesis lineage trajectories in mice (Fang et al., 2022).

**Figure 1:**
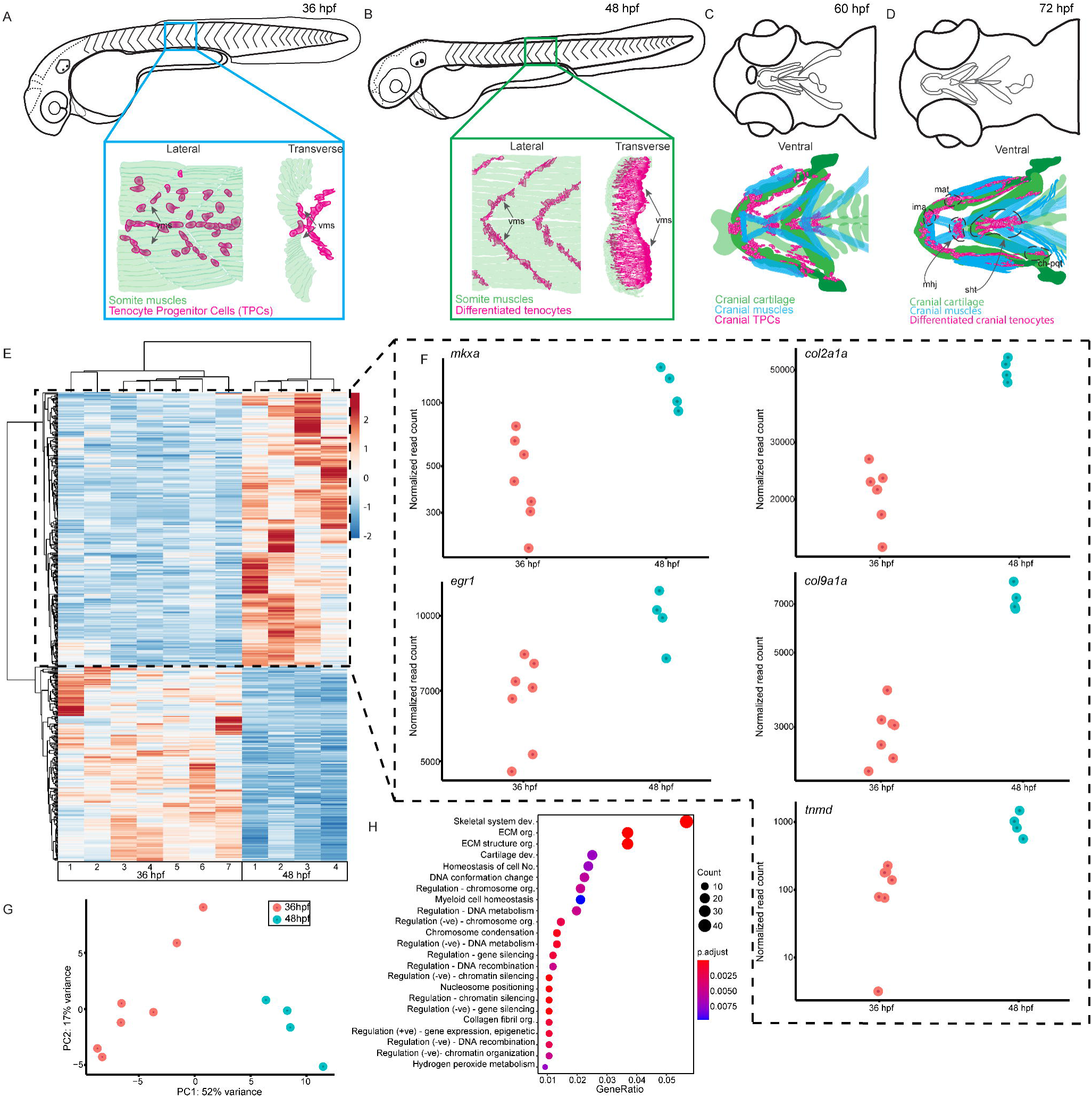
Onset of embryonic muscle contraction regulates transcription in tenocytes. **(A-D)** Diagrams depicting changes in tenocyte distribution and morphology during onset of trunk and cranial muscle contractions, **(A)** 36 hpf when twitching movements are sporadic, **(B)** 48 hpf when embryos become free swimming **(A-B)** Lateral views of 36 hpf **(A)** and 48 hpf embryos **(B)**. Insets show lateral and transverse views of migrating tenocyte progenitors **(A)** and differentiated tenocytes at somite boundaries with polarized, branched projections **(B)**. **(C-D)** Ventral views of the embryonic head in 60 **(C)** and 72 hpf **(D)** embryos at the onset of jaw movements. Cartilage (green), tenocytes (magenta) and muscles (cyan) showing tenocyte elongation, particularly in the sternohyoid tendon (sht) and condensation, particularly in the mandibulo-hyoid junction (mhj). **(E)** Heatmaps from bulk RNA-seq showing the top 1,000 differentially expressed genes (DEGs) between 36 hpf and 48 hpf. p < 0.05. **F)** Elevated expression of tenocyte marker genes *mkxa, tnmd,* and *egr1* and ECM genes *col2a1a*, *col9a1a* in RNA-seq experiments at 48 hpf. Datapoints represent normalized read counts of single biological replicates at each color-coded timepoint (n=7 for 36 hpf, n=4 for 48 hpf). **G)** Elevated expression of cartilage marker genes *col2a1a* and *col9a1a* in 48 hpf samples. **H)** PCA of individual replicates showing separation of experimental conditions by timepoint. **I)** GO analysis using Biological Process (BP) terms of top 2,788 DEGs by adjusted p-value.

To identify cell signaling pathways implicated in force-responses during embryonic tendon development, we analyzed our DEG list using ShinyGO (Ge et al., 2020) **(Supplementary File 2)** and DAVID **(Supplementary File 3)**, both of which interrogate KEGG pathway databases (Huang et al., 2009). ShinyGO identified DEGs associated with 52 different pathways with FDR < 0.05, including TGF-β, MAPK, Wnt, and Notch signaling, along with cell-cell adhesion and cell-ECM adhesion **(Supplementary File 2).** DAVID identified many of the same pathways as well as DEGs involved in RA metabolism, an emerging pathway of interest in tendon development (McGurk et al., 2017) **(Supplementary File 3)**.

Because our RNA-seq datasets were obtained from tenocytes during the onset of muscle contractions and swimming we also searched for DEGs associated with mechanosensitive pathways. Three genes of particular interest, *matn1*, *klf2a* and *mxra5b,* stood out based on their force-dependent regulation in other biological contexts or regulation by TGF- β, a well-known force-responsive signal (Maeda et al., 2011; Subramanian & Schilling, 2015). The top-most upregulated gene was *matn1,* which encodes an ECM protein highly enriched in cartilage; Matn1 enhances chondrogenesis of synovial fibroblasts treated with TGF-β (Pei et al., 2008). The transcription factor *klf2a* was also strongly upregulated; Klf2 and Klf4 have been implicated in enthesis development in mammalian tendons. Klf proteins also repress TGF-β signaling in endothelial cells (Boon et al., 2007; H. Li et al., 2021) and *klf2a* expression is mechanosensitive during heart valve development (Steed et al., 2016). The third DEG of particular interest was *mxra5b,* which encodes an ECM protein expressed in both tendons and ligaments during chick development (Robins & Capehart, 2018) and regulated by TGF-β in cultured human kidney epithelial cells (Poveda et al., 2017). Though other potentially mechanosensitive genes were present in our bulk RNAseq dataset, we focused on *matn1*, *klf2a* and *mxra5b* for further analysis based on evidence implicating them in mechanotransduction in other tissue contexts.

### *matn1*, *klf2a* and *mxra5b* are expressed in cranial and trunk tenocytes in vivo

To verify tenocyte-specific expression of *matn1*, *klf2a* and *mxra5b*, we performed in situ hybridization (ISH). Conventional chromogenic ISH for *matn1* detected no expression at 36 hpf but very strong expression at 48 and 60 hpf in developing craniofacial and pectoral fin cartilages **(Fig. 2—figure supplement 1A-C)**. Differential expression of *matn1* in our tendon dataset could reflect expression in developing fibrocartilage enthesis progenitors closely associated with cartilages. To test this idea, we conducted fluorescent in situ Hybridization Chain Reaction (*is*HCR) for *scxa* and *matn1* at 51 hpf, slightly later than our RNA-seq samples to allow better visualization of differentiated chondrocytes, and 72 hpf after the onset of jaw movements. *scxa*/*matn1* co-expressing cells localized to the intermandibularis anterior tendon (ima) and sternohyoid tendon (sht), specifically in the entheses that attach to meckels, anterior edge of the ceratohyal cartilages, and the posterior enthesis of the ceratohyal (ch-pqt), at 72 hpf **(Fig. 2A-F**, **Fig. 2—figure supplement 2A-D)**.

**Figure 2:**
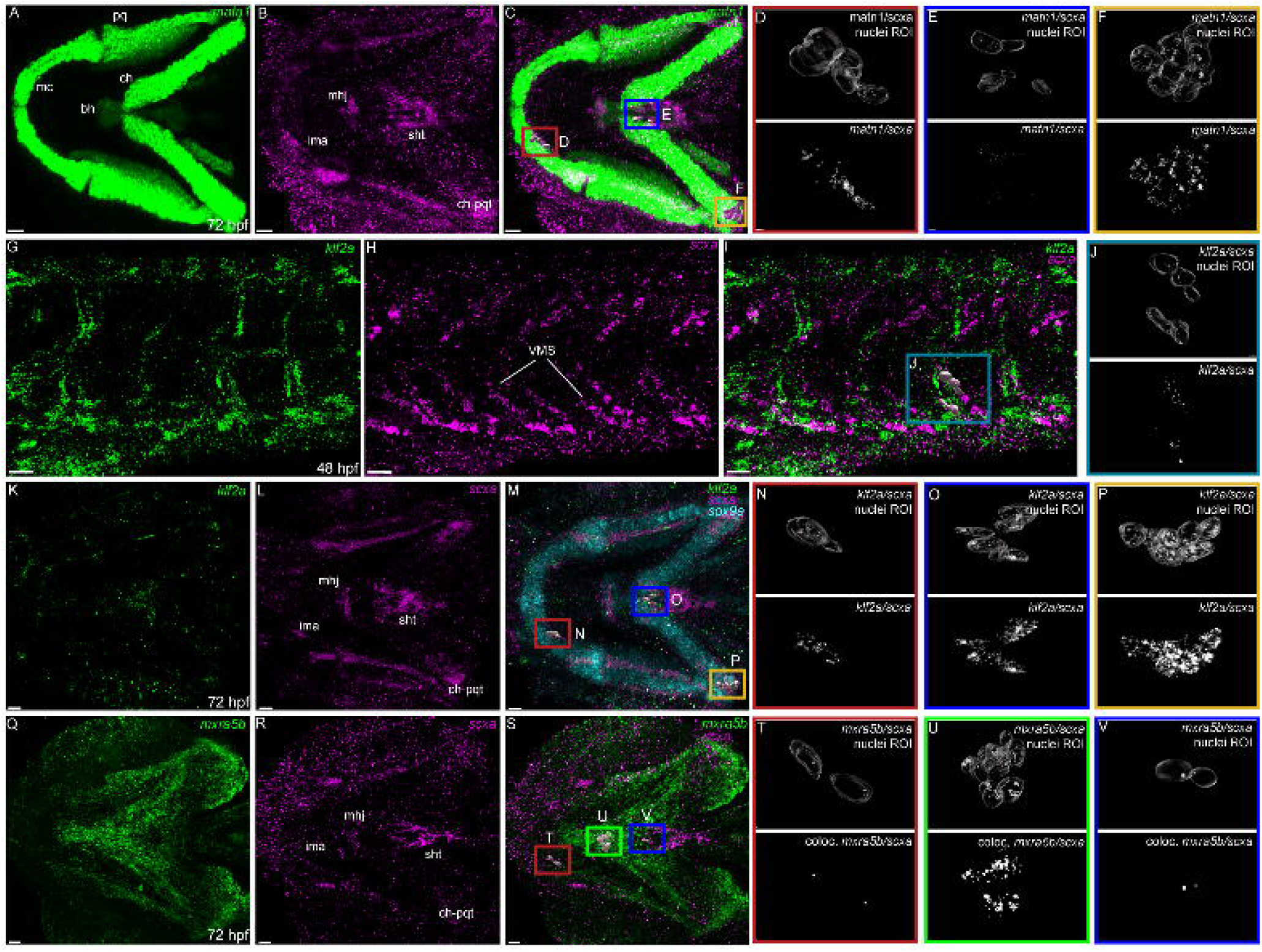
Expression of *matn1, klf2a* and *mxra5b* with *scxa* in cranial and trunk tenocytes. Ventral cranial **(A-F, K-V)** and lateral trunk **(G-J)** views of 72 hpf **(A-F, K-V)** and 48 hpf **(G-J)** embryos showing *is*HCR of *matn1* **(A,C,D-F)***, klf2a* **(G,I,J-K,M-P)**, and *mxra5b* **(Q,S-V)** in combination with *scxa* **(B-F,H-J,L-P,R-V)**. **(D-F, J, N-P, T-V)** Higher magnification views of tenocyte nuclei in marked ROI. **(C,D,M,N,S,T)** ROI and panels outlined in magenta show magnified views of 3D volumes of tenocytes associated with imt. **(I,J)** ROI and panels outlined in cyan show magnified views of 3D volume of vertical myoseptal (VMS) tenocytes. **(C,E,M,O,S,V)** ROI and panels outlined in royal blue show magnified views of 3D volume of tenocytes associated with sht enthesis. **(C,F,M,P)** ROI and panels outlined in yellow show magnified views of 3D volumes of tenocytes associated with ch-pqt. **(S,U)** ROI and panels outlined in green show magnified views of 3D volumes of tenocytes associated with mhj. Each magnified view of ROI displays a translucent outline of the nuclear 3D volume with white puncta representing voxel colocalizations of HCRish as depicted by the colocalization function in Imaris (see Methods).mc- Meckel’s cartilage, pq- palatoquadrate, ch-ceratohyal, bh- basihyal cartilage, ima- intermadibularis anterior tendon, mhj- mandibulohyoid junction, sht- sternohyoideus tendon, ch-pqt- ceratohyal-palatoquadrate tendon, sb-somite boundary. Scale bars = 20 microns.

For *klf2a,* chromogenic ISH revealed expression at vertical myosepta (VMS – somite boundaries) in the trunk at 48 hpf as well as developing pharyngeal arches and pectoral fins at 48 and 60 hpf **(Fig. 2—figure supplement 1D-F)**. This was confirmed by double *is*HCR of *klf2a* and *scxa* showing overlapping expression in tenocytes at VMS at 48 hpf **(Fig. 2G-J)**. *klf2a* expression was also detected in multiple cranial tendons at 72 hpf, most prominantly in the entheses of the ima, sht and ch-pqt **(Fig. 2K-P)**. This provides the first evidence for *klf2a* as an enthesis marker in craniofacial tendons, similar to Klf2 expression in developing mouse limb entheses (Kult et al., 2021; Lu & Thomopoulos, 2013; Zelzer et al., 2014).

*mxra5b* expression was first detected by chromogenic ISH at VMS near the horizontal myoseptum (HMS), which separates dorsal and ventral somites at 36 hpf, as well as in the notochord and cranial mesenchyme at 48 hpf **(Fig. 2—figure supplement 1G-H)**. Expression increased and extended along the VMS by 60 hpf **(Fig. 2—figure supplement I)**. Double isHCR for *scxa* and *mxra5b,* detected *mxra5b* expression in cranial entheses (including ima, mhj, and sht – as well as others not shown), and in the mandibulohyoid junction tendon (mhj) in embryos at 72 hpf **(Fig. 2Q-V)**. Similar to *klf2a*, *mxra5b* expression has not been described in cranial connective tissues previously.

### Tenocyte-specific gene expression of *matn1*, *klf2a* and *mxra5b* is regulated by muscle contraction

Since *matn1, klf2a* and *mxra5b* were among the top DEGs in tenocytes at the onset of active swimming and persistent muscle activity, we reasoned that mechanical force regulates their expression. To test this, we performed Real Time Quantitative-PCR (RT-qPCR) in genetically paralyzed embryos. Relative expression of each gene was compared between wild-type (WT) embryos and homozygous mutants lacking the function of the voltage dependent L-type calcium channel subtype beta-1 (*cacnb1^-/-^*), which blocks muscle contraction (Subramanian et al., 2018; Zhou et al., 2006). At 48 hpf, all 3 genes were downregulated in *cacnb1^-/-^*mutants versus WT **(Fig. 3—figure supplement 1A)**. In contrast, at 72 hpf once jaw movements had begun, only *matn1* and *mxra5b* remained downregulated in *cacnb1^-/-^* embryos while *klf2a* expression increased **(Fig. 3—figure supplement 1B)**.

**Figure 3:**
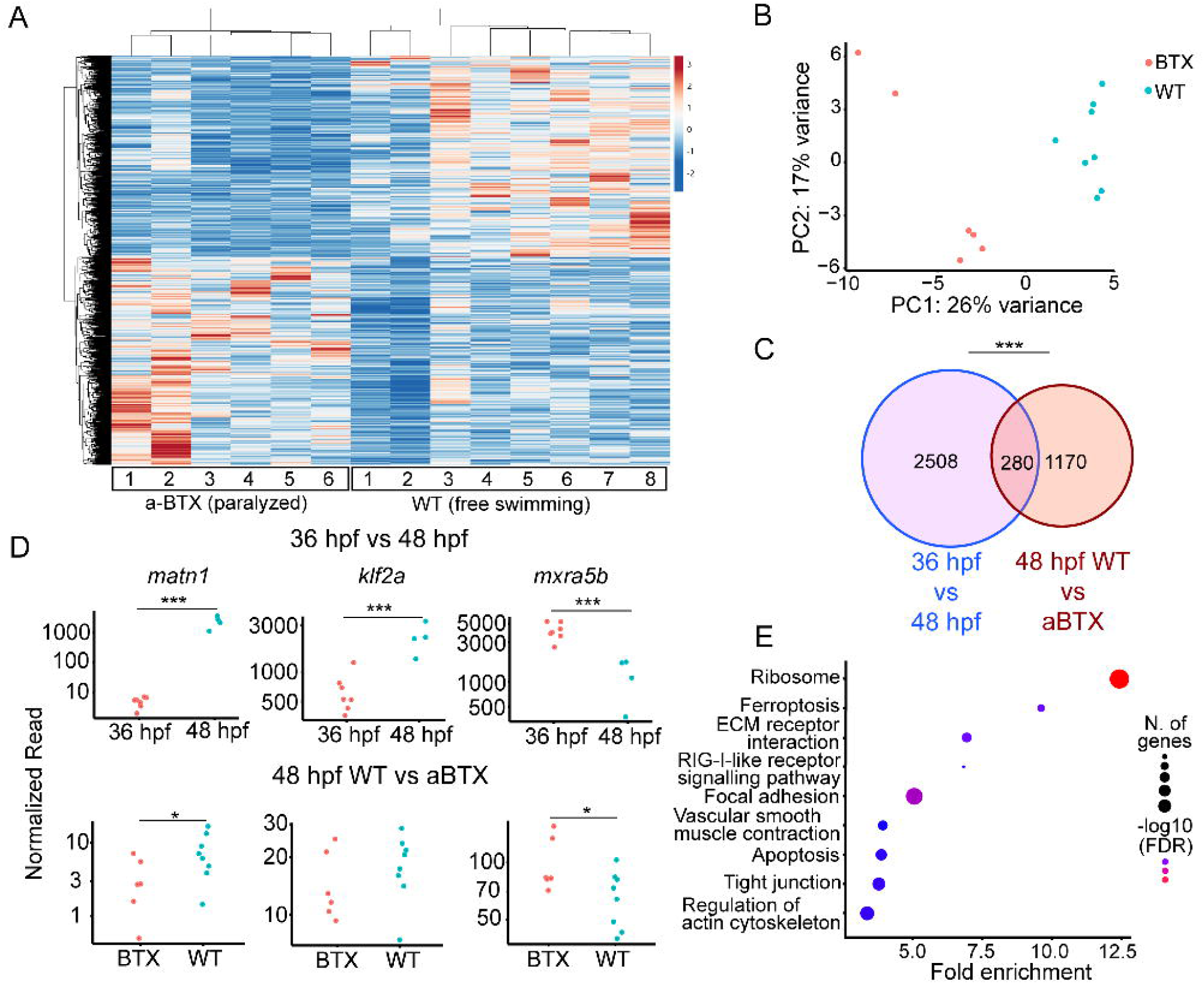
Paralysis regulates tenocyte gene expression in developing musculoskeletal system. **(A)** Heatmap of differentially expressed genes (DEG) from bulk RNA-seq between WT and *αBTX*-injected (*αBTX*-inj) paralyzed 48 hpf embryos (force perturbed). **(B)** PCA of individual replicates WT versus *αBTX*-inj embryos’ RNA-seq separate by experimental condition. **(C)** Venn diagram shows overlap of genes between developmental time-point and force perturbed RNA- seq experiments. **(D)** Comparison of normalized read counts between replicates of *matn1, klf2a,* and *mxra5b* in 36 hpf vs. 48 hpf and WT versus *αBTX* RNA-seq experiments. **(E)** KEGG pathway analysis plot shows enrichment of overlapping genes from **(C)**. ns = not significant, * p < 0.05, ** p < 0.01, *** p < 0.001

To confirm that loss of muscle contraction caused these transcriptional changes in tenocytes we injected *Tg(scxa:mCherry)* embryos at the 1-cell stage with mRNA encoding full-length alpha-bungarotoxin mRNA (αBTX), which paralyzes embryos by irreversibly binding to acetylcholine receptors at neuromuscular synapses. Bulk RNA-seq of sorted *mCherry+* cells from whole αBTX-injected embryos at 48 hpf compared with WT uninjected controls **(Fig. 3A)** identified 1,450 DEGs. PC analysis clearly separated WT and αBTX biological replicates **(Supplementary File 4)** (**Fig. 3B)**. 280 DEGs overlapped between both bulk RNA-seq runs **(Fig. 3C**, **Supplementary File 5)**. KEGG pathway analysis, using shinyGO (Ge et al., 2020)identified many of the same pathways downregulated in *cacnb1^-/-^*embryos, as well as others not previously implicated in tendon mechanotransduction. Seveal of these mapped to terms such as “Focal Adhesion”, including *rhoab, rock2a* (both part of Rho-ROCK signaling), and *col9a1a* **(Supplementary File 6)** (**Fig. 3E)** further implicating these as force-dependent in tendons.

Comparisons of *matn1, klf2a,* and *mxra5b* expression between αBTX and WT versus our original 36 hpf versus free swimming 48 hpf RNA-seq experiment, revealed similar trends in expression. This suggests that the expression changes seen at embryonic stages (36 hpf vs 48 hpf) reflect tenocyte responsiveness to muscle contraction **(Fig. 3D)**. Further, comparing the 48 hpf WT versus *cacnb1^-/-^* mutant RT-qPCR with both bulk RNAseq experiments, *matn1* and *mxra5b* expression were both consistently downregulated by paralysis, while *klf2a* expression was more variable across experiments **(Fig. 3A**, **3D)**.

Having shown reproducible changes in their expression between bulk RNA-seq results, we next asked if variable recovery of muscle contractile forces differentially affects changes in *matn1, klf2a,* and *mxra5b* expression caused by paralysis. To test this, we used 90ng/ul of full-length aBTX mRNA, a concentration optimized to paralyze embryos only for the first two days of embryogenesis after which they gradually recover. Nearly all aBTX-injected embryos regained muscle contractions and were swimming at 48 hpf. We performed RT-qPCR on cDNA derived from these embryos and compared them to aBTX paralyzed (aBTX-P) and uninjected controls. We separated 48 hpf recovered embryos into two subgroups based on the extent of muscle contraction: 1) partially recovered (Twitching or aBTX-T), in which embryos showed sporadic contractions of the trunk and pectoral fin muscles, similar to twitching 36 hpf embryos and 2) fully recovered (Recovered, or aBTX-R), in which embryos swam freely. At 48 hpf, RT-qPCR revealed significant global downregulation of *matn1* and *mxra5b* in αBTX paralyzed embryos compared to WT uninjected siblings, like *cacnb1^-/-^* mutant embryos **(Fig. 3—figure supplement 1C)** and were upregulated in aBTX-T and aBTX-R embryos **(Fig. 3—figure supplement 1D)**. In contrast, *klf2a* was upregulated in paralyzed embryos, though this increase was also not statistically significant from WT controls **(Fig. 3—figure supplement 1E)**. These results, combined with those from RNA-seq, suggest that *matn1, klf2a,* and *mxra5b* transcription during development are regulated by muscle contraction.

To verify that these transcriptional changes occur specifically in tenocytes in response to force, we examined *matn1, klf2a*, and *mxra5b* expression in *scxa*-positive cells by *is*HCR with mCherry antibody staining of *Tg(scxa:mCherry)*) fish using our αBTX paralysis-recovery experimental protocol **(Figs. 4-6)**. Additionally, we quantified expression at multiple attachment regions across different tendons for each gene to determine if responses differed between spatially distinct tendons and by attachment type (e.g. enthesis versus MTJ).

**Figure 4:**
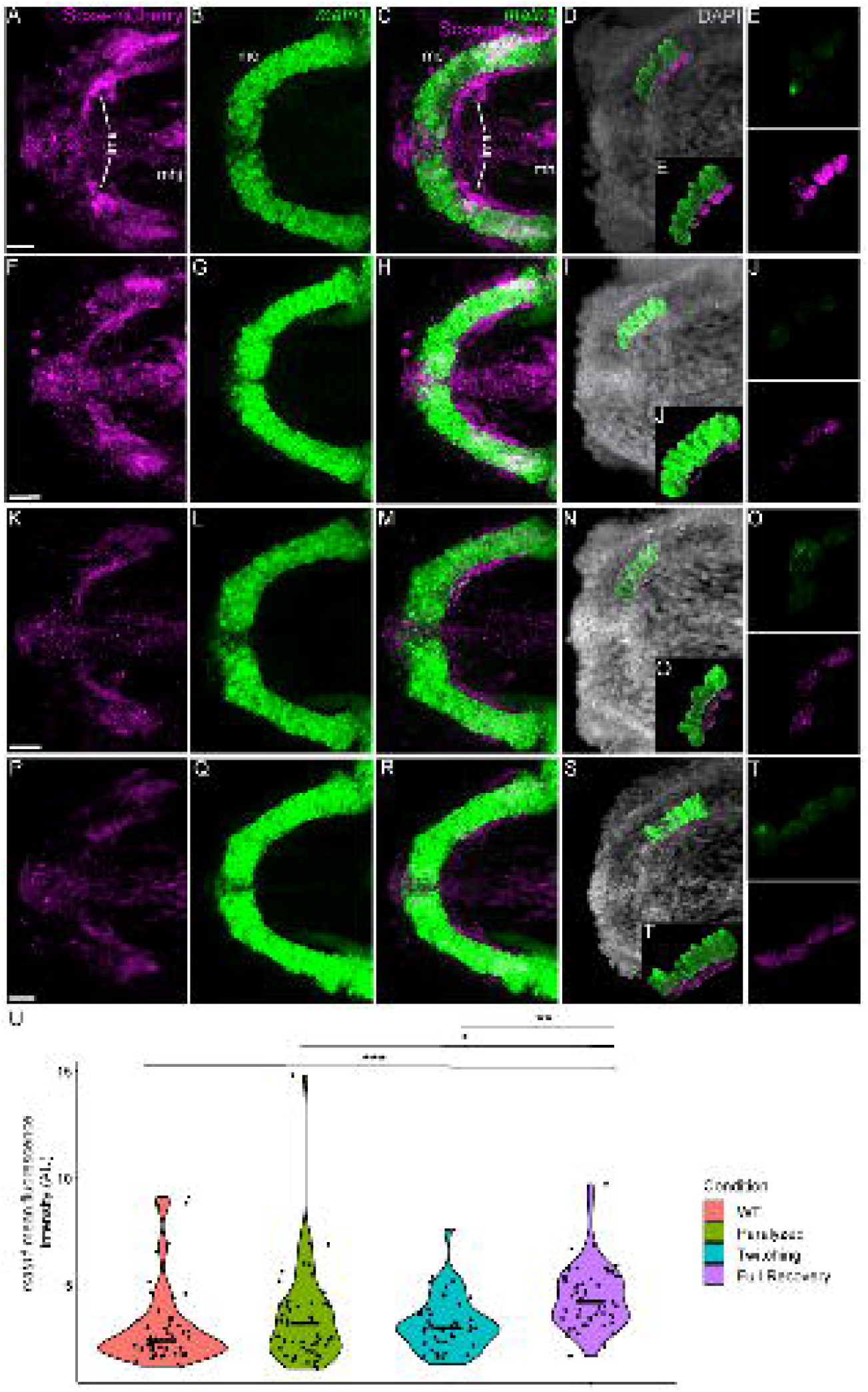
Mechanical force differentially regulates expression of *matn1* in ima enthesis tenocytes. Ventral views of Meckel’s cartilage and associated tenocytes showing *is*HCR of *matn1* (green) and anti-mCherry immunofluorescence (magenta) marking the tenocytes in *Tg(scxa:mCherry)* embryos at 72 hpf in WT uninjected (WT) **(A-E)**, *αBTX*-inj paralyzed **(F-J)**, partially recovered *αBTX*-inj (Twitching) **(K-O)**, and completely recovered *αBTX*-inj (Full Recovery) **(P-T)** conditions at ima enthesis. **(D,I,N,S)** Grayscale images showing nuclei stained with DAPI with ROIs showing isolated 3D-volumes of chondrocytes (green) and enthesis tenocytes (magenta) based on DAPI signal. **(E,J,O,T)** Insets showing magnified views of the 3D-volumes of tenocytes associated with ima enthesis depicting expression of *matn1* and stained for mCherry. **(U)** Violin plot showing changes in mean fluorescence intensity of *matn1* in ima enthesis tenocyte nuclei between WT (n=8), Paralyzed (n=8), Twitching (n=6) and Full Recovery (n=7) with ∼8 nuclei measured per embryo. p-value calculated with linear mixed effects model with Tukey post-hoc test. * p < 0.05, ** p < 0.01, *** p < 0.001). Scale bars = 20 microns. Figure 4—source data 1 Measurements of *matn1* HCR signal intensity in ima enthesis tenocytes [Figure 4 Source Data.xlsx]

For *matn1*, we quantified expression by measuring its fluorescence intensity in individual tenocytes in 3D at the intermandibularis anterior (ima) enthesis where the ima attaches to meckel’s (mc) cartilage and the sht enthesis at the anterior end of the ch cartilage **(Fig. 4A-T**, **Fig. 4—figure supplement 1A-L)** (Subramanian et al., 2023). Cells were selected for quantification by their co-expression of *matn1* and *scxa* and positions near chondrocytes expressing *matn1* alone and tenocytes expressing *scxa* alone, as described previously (Subramanian et al., 2023). In these ima tenocytes, we found no significant difference in *matn1* expression between WT and paralyzed embryos, but increased expression in fully recovered (aBTX-R) embryos relative to WT, Paralyzed, and Twitching (aBTX-T) embryos **(Fig. 4U)**. Conversely, tenocytes of the sht enthesis showed no significant difference in expression across any of the conditions **(Fig. 4—figure supplement 1M)**.

We also examined fluorescence intensity in *scxa/mxra5b* or *scxa/klf2a* double positive tenocytes located at ima and sht entheses, as well as mhj and sht MTJs. m*xra5b* expression in the ima enthesis was significantly reduced in paralyzed, aBTX-T twitching, and remained low in aBTX-R fully recovered embryos compared to WT **(Fig. 5—figure supplement 1Z)**. However, in tenocytes of all other measured attachment sites (sht enthesis, mhj MTJ, sht MTJ), *mxra5b* expression returned to WT levels upon full recovery **(Fig. 5**, **Fig. 5—figure supplement 2M, Fig. 5—figure supplement 3M)**. *klf2a* expression in ima and sht entheses was significantly increased in paralyzed and aBTX-T embryos compared to WT and further increased upon full recovery **(Fig. 4—figure supplement 1Z, Fig. 5—figure supplement 1M)**. However, unlike entheses, klf2a expression in sht MTJ tenocytes only increased significantly from twitching to full recovery, and in mhj MTJ tenocytes the pattern was much more variable, increasing upon paralysis, decreasing to WT levels at twitching, and re-increasing beyond WT levels at full recovery **(Fig. 5—figure supplement 3, Fig. 6U)**.

**Figure 5:**
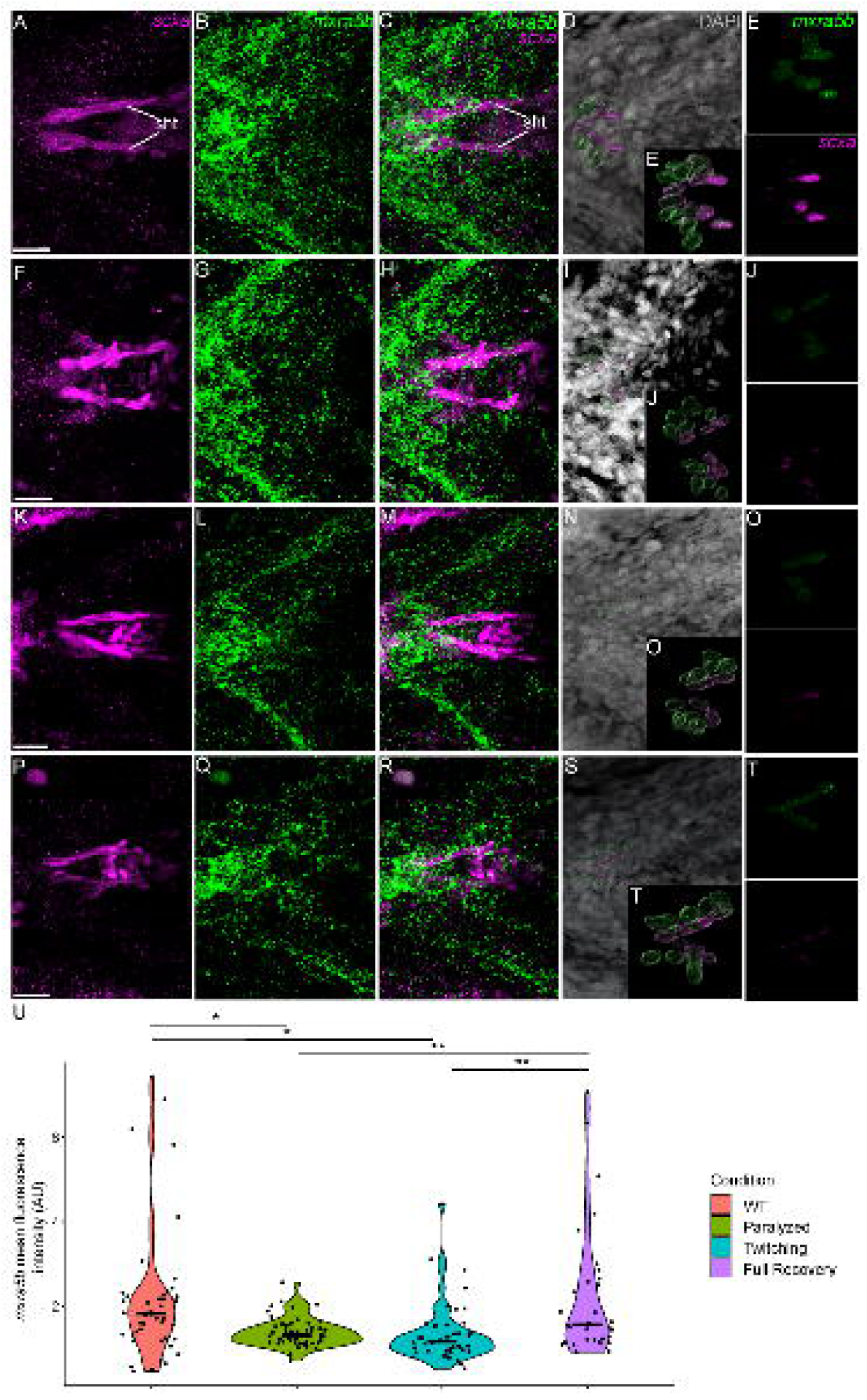
Mechanical force differentially regulates expression of *mxra5b* in sh-ch (sht) enthesis tenocytes. Ventral views of ceratohyal (ch) cartilage and associated tenocytes showing *is*HCR of *mxra5b* (green) and anti-mCherry immunofluorescence (magenta) marking the tenocytes in *Tg(scxa:mCherry)* embryos at 72 hpf in WT uninjected (WT) **(A-E)**, *αBTX*-inj paralyzed **(F-J)**, partially recovered *αBTX*-inj (Twitching) **(K-O)**, and completely recovered *αBTX*-inj (Full Recovery) (P-T) conditions at ima enthesis. **(D,I,N,S)** Grayscale images showing nuclei stained with DAPI with ROIs showing isolated 3D-volumes of chondrocytes (green) and sternohyoideus- ceratohyal (sh-ch) enthesis tenocytes (magenta) based on DAPI signal. **(E,J,O,T)** Insets showing magnified views of the 3D-volumes of tenocytes associated with sh-ch enthesis depicting expression of *mxra5b* and stained for mCherry. **(U)** Violin plot showing changes in mean fluorescence intensity of *mxra5b* in sh-ch enthesis tenocyte nuclei between WT (n=7), paralyzed (n=8), twitching (n=8) and full Recovery (n=4) with ∼8 nuclei measured per embryo. p-value calculated with linear mixed effects model with Tukey post-hoc test. * p < 0.05, ** p < 0.01, *** p < 0.001). Scale bars = 20 microns. Figure 5—source data 3 Measurements of *mxra5b* HCR signal intensity in sht enthesis tenocytes [Figure 5 Source Data.xlsx]

**Figure 6:**
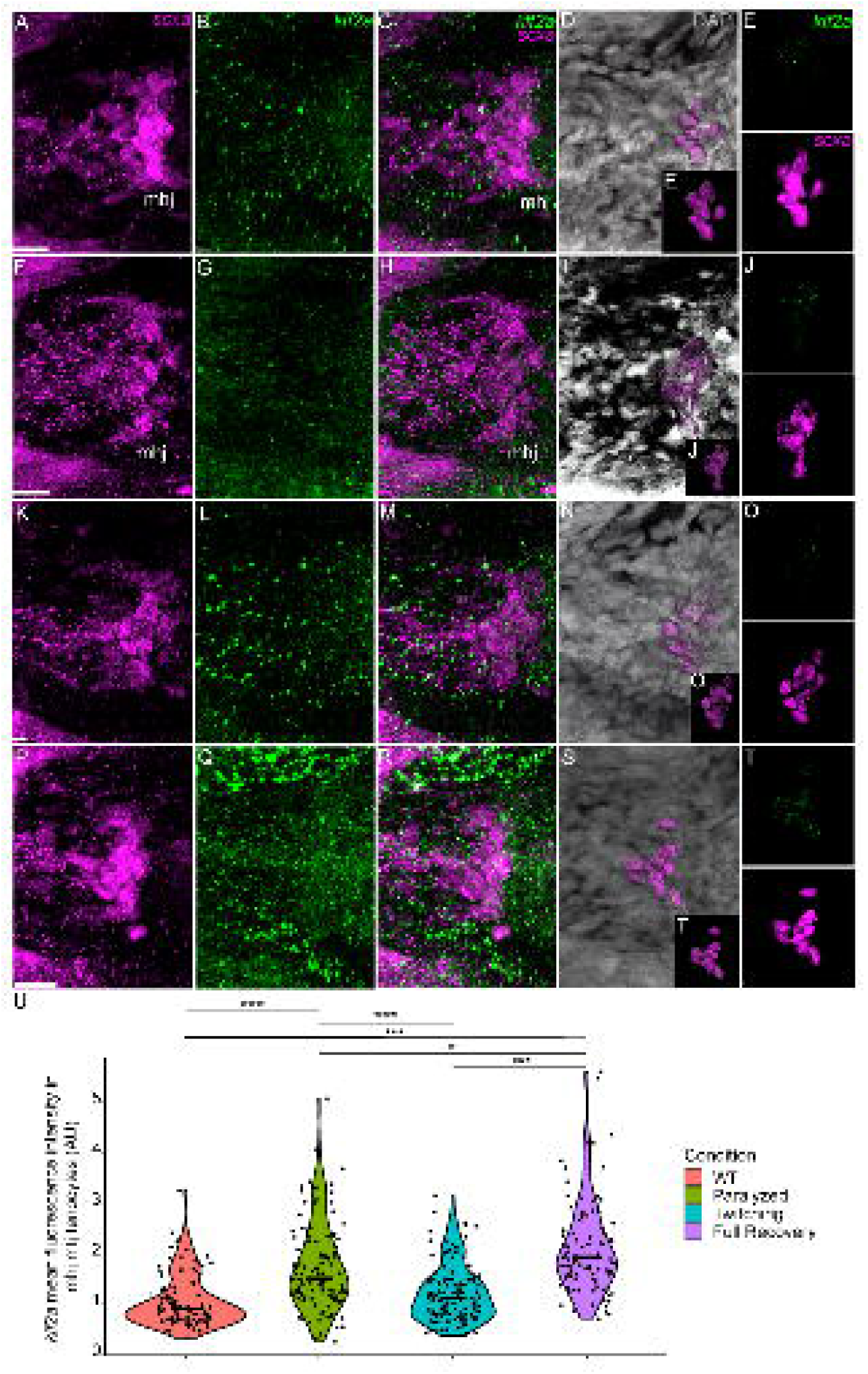
Mechanical force regulates expression of *klf2a* in mhj myotendinous tenocytes. Ventral views of mandibulohyoid(mhj) myotendinous junction (mtj) associated tenocytes showing *is*HCR of *klf2a* (green) and anti-mCherry immunofluorescence (magenta) marking the tenocytes in *Tg(scxa:mCherry)* embryos at 72 hpf in WT uninjected (WT) **(A-E)**, *αBTX*-inj paralyzed **(F-J)**, partially recovered *αBTX*-inj (Twitching) **(K-O)**, and completely recovered *αBTX*-inj (Full Recovery) **(P-T)** conditions. **(D,I,N,S)** Grayscale images showing nuclei stained with DAPI with ROIs showing isolated 3D-volumes of mhj tenocytes (magenta) based on DAPI signal. **(E,J,O,T)** Insets showing magnified views of the 3D-volumes of tenocytes associated with mhj-mtj depicting expression of *klf2a* and stained for mCherry. **(U)** Violin plot showing changes in mean fluorescence intensity of *klf2a* in mhj-mtj tenocyte nuclei between WT (n=17), Paralyzed (n=15), Twitching (n=14) and Full Recovery (n=11) with ∼10 nuclei measured per embryo. p-value calculated with linear mixed effects model with Tukey post-hoc test. * p < 0.05, ** p < 0.01, *** p < 0.001). Scale bars = 20 microns. Figure 6 source data 7 Measurements of *klf2a* HCR signal intensity in mhj mtj tenocytes [Figure 6 Source Data.xlsx]

To address functions of *matn1, klf2a,* and *mxr5b* in tenocytes we used multiplex CRISPR/Cas9 mutagenesis (R. S. Wu et al., 2018) to generate F0 crispants for *matn1*, *klf2a* and *mxra5b*. While we did not observe obvious phenotypic defects in *matn1* and *klf2a* crispants, possibly due to genetic redundancy with other similar proteins, *Tg(scxa:mCherry)* embryos injected with 4 *mxra5b* gRNAs had qualitatively fewer trunk tenocytes when compared to uninjected controls **(Fig. 7A)**. Additionally, trunk VMS in *mxra5b* CRISPR-injected embryos displayed a wider sb angle **(Fig. 7B)**, although this may reflect a role for mxra5b in the notochord, where it is also expressed **(Fig. 2—figure supplement 2G-I)**. These results suggest that mxra5b may be required for embryonic axial tenocyte migration and/or differentiation.

**Figure 7:**
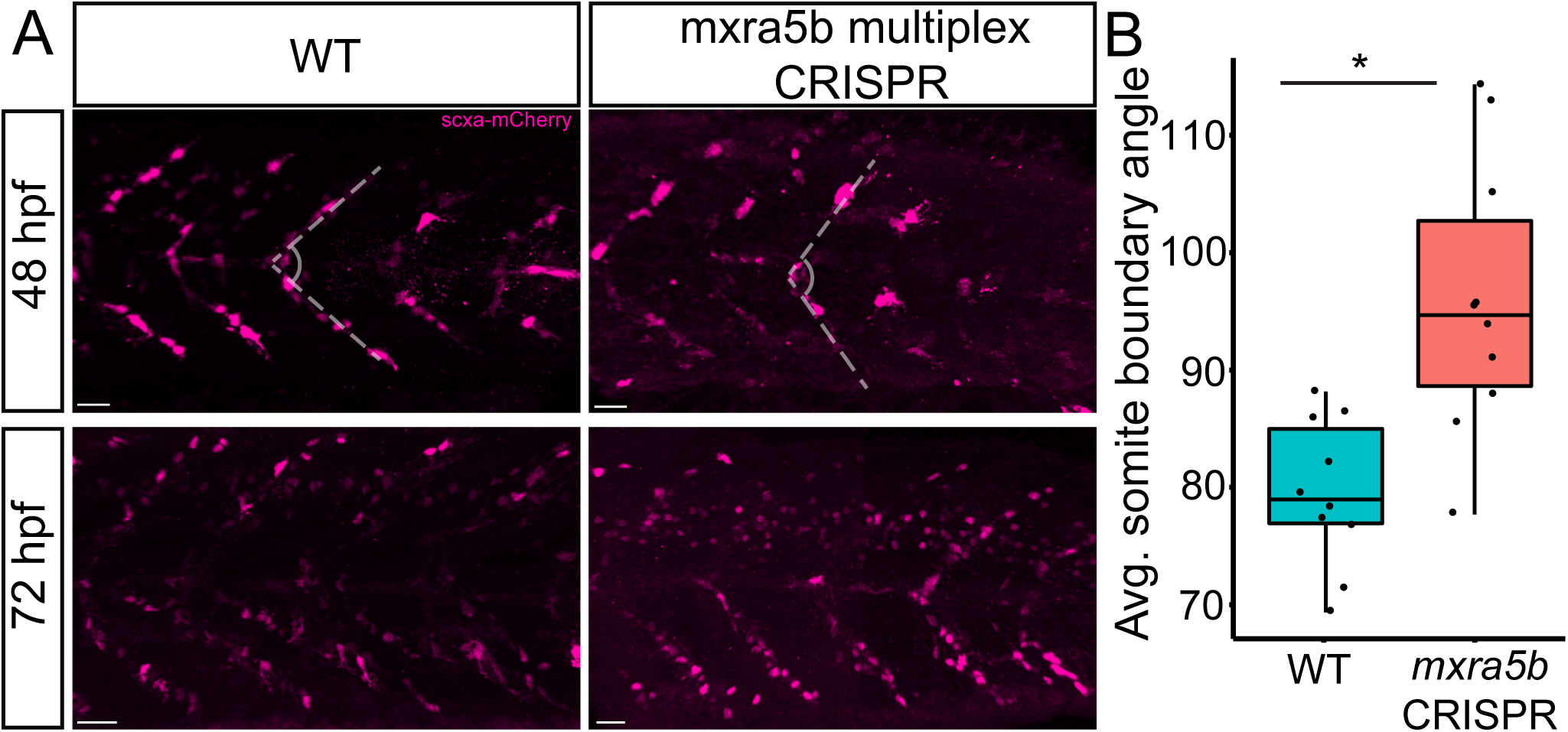
Loss of mxra5b function affects somite boundary structure. **(A)** Lateral views of WT and *mxra5b* multiplex CRISPRants at 48 hpf and 72 hpf Tg(*scx:mCherry*) embryos stained with anti-mCherry to show tenocytes at the somite boundary (SB). **(B)** Quantification of somite boundary angle measurements of 48 hpf WT or *mxra5b* multiplex CRISPRant embryos. P-value calculated with Watson’s U2 test. * p < 0.05

## Discussion

Previous studies of mechanotransduction in tenocytes, particularly at the transcriptional level, have largely been limited to adult tendons or in vitro assays using mesenchymal stem cells. Few have addressed how functional differences in tendons are established during embryonic development. We report the first genome-wide survey of embryonic mechanoresponsive genes and their transcriptional responses across multiple tendon types. and identify three genes induced at the onset of muscle attraction and later maintained by contractile force **(Fig. 8A)**. Paralysis of zebrafish embryos alters expression of two ECM proteins in tenocytes, *matn1* and *mxra5b*, as well as the transcription factor *klf2a*. All three are expressed in cranial entheses, while *mxra5b* and *klf2a* are also expressed in trunk MTJs **(Fig. 2**, **Fig. 2—figure supplement 1, Fig. 2—figure supplement 2, Fig. 8B)**. Our previous studies have shown that in both tissues embryonic tenocytes in zebrafish acquire specialized morphologies and gene expression profiles as muscles first form functional attachments (Subramanian et al., 2018, 2023; Subramanian & Schilling, 2015). In contrast to classical studies of mature tendons these results suggest that cells with distinct enthesis or MTJ signatures arise in the embryo to fine-tune the ECM to match the functional demands of and forces exerted by individual muscles.

**Figure 8:**
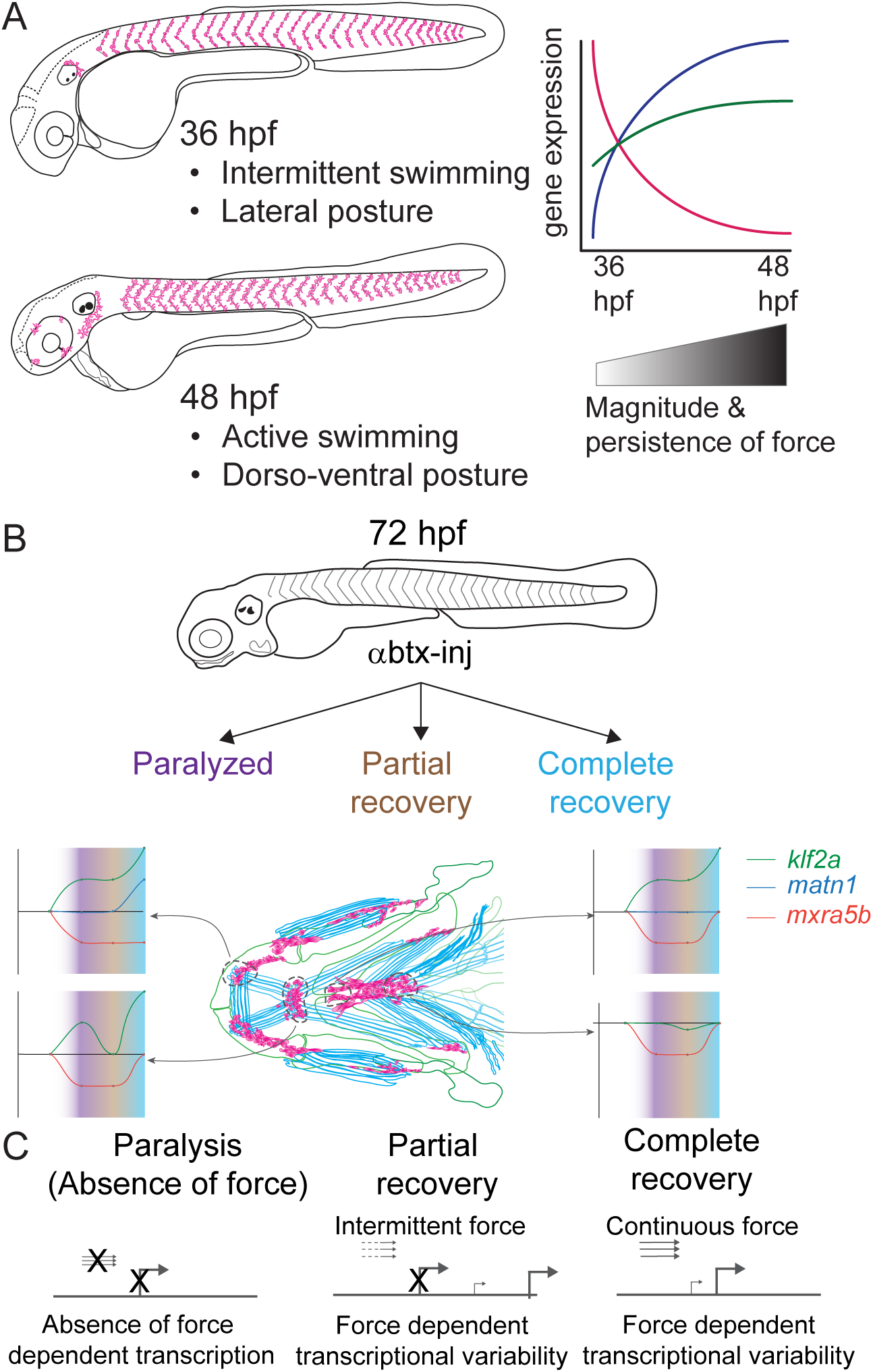
Model depicting role of mechanical force in regulating expression of genes in tenocytes during onset of active muscle contraction. **A)** Cartoon showing role of force in regulating tenocyte morphogenesis and gene expression in tenocytes between 36 hpf and 48 hpf stages correlating with onset of active swimming The variability in gene expression is related to increase in both magnitude and persistence of muscle contraction force. **B)** Representative model summarizing the multifaceted role of muscle contractile force on expression dynamics of *matn1, klf2a*, and *mxra5b* genes in cranial tendon attachments. **C)** Force-responsive gene expression is more nuanced than a binary on/off control.

Classically tendon types and subdomains are distinguished by their collagen composition, and many collagens are direct Scx or Mkx transcriptional targets (Bobzin et al., 2021; Felsenthal & Zelzer, 2017; Subramanian & Schilling, 2015). This helps explain the gradient of stiffness and corresponding Scx/Sox9 expression within an enthesis (Blitz et al., 2013; Lu & Thomopoulos, 2013; Subramanian et al., 2023; Zelzer et al., 2014). Our results highlight additional genes implicated in cartilage (i.e. *matn1*) and fibrocartilage (i.e. *KLF*) in entheseal tenocytes and their force responses. Though typically thought of as cartilage-specific, *matn1* and its relatives have been reported in single-cell RNA-seq (scRNA-seq) analyses of adult tenocytes and fibrocartilage (Kaji et al., 2021). We find that zebrafish *matn1* regulation differs between entheses that form at different stages **(Fig. 4**, **Fig 4—figure supplement 1)**. Whereas paralyzed embryos at both twitching and swimming stages show reduced tenocyte *matn1* expression **(Fig. 3D)**, our HCRish data reveal that expression only rebounds after full recovery of muscle contraction in the ima enthesis **(Fig. 8B**) **(Fig. 4**, **Fig. 4—figure supplement 1)**. These spatial and temporal differences support our hypothesis that these are bona fide embryonic entheseal tenocytes specified at the edges of cartilages as muscles first attach (Subramanian et al., 2023). They are also consistent with studies showing that *matn1* transcription is upregulated upon mechanical load in cultured chondrocytes (Chen et al., 2016). Chondrocyte ECM becomes disorganized in *Matn1^-/-^* mutant mice exposed to mechanical loads after medial meniscus destabilization surgery (Chen et al., 2016; P. Li et al., 2020). Our data implicate *matn1* in tendon/fibrocartilage mechanotransduction and in the initial establishment of ECM stiffness gradients at entheses during embryogenesis **(Fig. 4**, **Fig. 4—figure supplement 1)** (Lu & Thomopoulos, 2013).

*Mxra5* (also known as *adlican*) encodes a secreted proteoglycan implicated in cell-cell adhesion and ECM remodeling, mainly in the context of colorectal and other cancers (He et al., 2015; G.-H. Wang et al., 2013). *Mxra5* is expressed in tendons and other connective tissues of developing chick embryos as well as human fibroblasts (Chondrogianni et al., 2004; Robins & Capehart, 2018). We find that zebrafish *mxra5b* expression is downregulated in all tenocytes at the onset of embryonic muscle contraction, unlike *matn1* **(Fig. 3**, **Fig. 5**, **Fig. 5—figure supplements 1, 2, 3)**. Consistent with a force-responsive gene, MXRA5 is inhibited by TGF-β1 (Poveda et al., 2017), and associated with migration of dental pulp stem cells (Yoshida et al., 2023). Our results provide the first evidence for regulation of *mxra5b* transcription in tenocytes by mechanotransduction. However, despite reductions in *mxra5b* levels overall with loss of active muscle contraction, our HCRish results suggest that these changes differ between distinct tendons and force conditions **(Fig. 5)**. For example, in the ima enthesis, paralysis downregulates *mxra5b* expression, with little rebound after recovery **(Fig. 8B**) **(Fig. 5—figure supplement 1)**. In contrast, at other entheses and MTJs *mxra5b* expression returns to WT levels upon full recovery after paralysis **(Fig. 8B**) **(Fig. 5**, **Fig. 5—figure supplements 2 & 3)**. *mxra5b* expression may require continuous mechanical activation, levels of which differ between tendons as well as entheses or MTJs **(Fig. 8B)**. This heterogeneity may help explain differences between our RNAseq results for *mxra5b* and *is*HCR expression data, since the RNAseq experiments were performed on FAC sorted tenocytes of all tendons **(Fig. 3)**.

Similar to *matn1* and *mxra5b,* 1) zebrafish *klf2a* expression localizes to embryonic cranial entheses, 2) its transcription increases in in tenocytes at the onset of muscle contraction, and 3) these responses vary between spatially distinct tendons and tendon subdomains **(Fig. 8B**) **(Fig. 3**, **Fig. 4—figure supplement 1, Fig. 5—figure supplements 1 & 3**, **Fig. 6)**. Mammalian Klf2 and Klf4 have been implicated in cell differentiation at tendon-bone entheses (Kult et al., 2021). Cranial tenocytes in zebrafish upregulate *klf2a* upon recovery from paralysis **(Fig. 3**, **Fig. 4—figure supplement 1, Fig. 5—figure supplements 1 & 3, Fig. 6)**, though there are discrepancies between HCRish, bulk RNAseq, and qPCR measurements. These may reflect the fact that *klf2a* is also expressed in other tissues, such as embryonic vascular and endocardial cells **(Fig. 2)** (Goddard et al., 2017) or differences in expression between trunk and cranial tenocyte populations. The HCRish data show distinct entheseal *klf2a* and MTJ expression patterns **(Fig. 4—figure supplement 1, Fig. 5—figure supplements 1 & 3, Fig. 6) (Fig. 8B)**. Klf2 binding sites have been identified upstream of ECM genes such as *Col5* in sorted entheseal tenocytes (Kult et al., 2021). *Klf2* expression is also upregulated by fluid forces in endocardial cells leading to fibronectin synthesis (Boselli et al., 2015; Lee et al., 2006; Steed et al., 2016). Thus, force dependent *klf2a* expression may be critical for tissue-specific ECM remodeling in many contexts.

Together, our bulk RNA-seq analysis of embryonic zebrafish tenocytes and their transcriptional responses to muscle contraction: 1) identifies new regulators of tenocyte-ECM, going beyond the better studied collagens, and 2) highlights the importance of considering developmental events that specify the mechanical properties of tendons as they form. Genes such as *matn1, mxra5b* and *klf2a* show unique expression profiles and changes due to perturbation of muscle contraction, both during normal embryonic development and in response to paralysis **(Fig. 8B)**. The presence of these genes in embryonic tendons and their responses to force during normal development versus recovery from paralysis raises questions as to whether the mechanisms that initially establish these structures differ from those that control their maintenance **(Fig. 8B)**. Though cell-ECM feedback mechanisms have been studied in controlled 3D microenvironments in-vitro, extrapolating these mechanisms into an understanding of in vivo biological processes like development and tissue homeostasis is necessary (Saraswathibhatla et al., 2023). Given the large variation of cell-ECM feedback mechanisms throughout embryonic development, understanding specific tenocyte-ECM interactions will require novel approaches to measuring the effect of varying 1) ECM microenvironment protein compositions, or local “matrisomes,” on tenocyte gene expression and 2) intrinsic gene expression patterns of heterogeneous tenocyte populations spatially and functionally. Single-cell approaches (e.g. scRNA-seq) at different developmental stages and in the presence or absence of force, will provide a clearer understanding of how individual spatially and functionally distinct tenocyte populations respond to force in development. Integrating such knowledge of the basic biology of tenocytes at multiple scales will be essential for developing a better picture of tenocyte-ECM interactions at individual tendons, paving the path to advance personalized translational therapies for tendon injuries.

## Materials and Methods

### Zebrafish embryos, transgenics and mutants

Wild type zebrafish ( AB strain), *TgBAC(scxa:mCherry)^fb301^*transgenics referred to as *Tg(scxa:mCherry)*), or *cacnb1^ir1092/ir109^;fb301Tg* (referred to as *cacnb1^-/-^* mutants) embryos were raised in embryo medium at 28.5°C (Westerfield, 2000) and staged as described (Kimmel et al., 1995). Craniofacial musculoskeletal structures were identified and annotated as described previously (Schilling & Kimmel, 1997; Subramanian et al., 2023). All protocols performed on embryos and adult zebrafish in this study had prior approval from the IACUC at UC Irvine.

### In situ hybridization (ISH)

Digoxigenin labelled antisense RNA probes for *matn1*, *klf2a*, and *mxra5b* were generated using T7 sequence-tagged primers **(Supplementary File 7)**. Total embryo RNA was extracted from 72 hpf WT embryos using Trizol (Invitrogen 15596026) and Monarch Total RNA Miniprep kit (New England Biolabs (NEB) T2010S). cDNA was synthesized using olido dT primers from ProtoScript II First Strand cDNA Synthesis Kit (NEB E6560) and used as a template to synthesize RNA probes using T7 RNA polymerase (Roche, 10881767001) and DIG RNA labelling mix (Roche, 11277073910). Whole-mount ISH was performed with anti-DIG-AP fragments (Roche, 11093274910) at 1:2000 dilution, as described in (Thisse & Thisse, 2008).

### In situ hybridization chain reaction (HCRish) and immunohistochemistry

HCRish probes were designed by Molecular Instruments Inc. (Los Angeles, CA) and whole mount HCRish was performed with amplifiers/probes obtained from Molecular Instruments according to the HCRish v3.0 protocol as described (Choi et al., 2014; Subramanian et al., 2023; Trivedi et al., 2018). Probes/amplifier combinations used were: *matn1* (NCBI ref. # 403023) and *mxra5b* (NCBI ref. # 795448) in B1 with B1 Alexa Fluor 488, *scxa* (NCBI ref. # 100034489) in B2 with B2 Alexa Fluor 546, *klf2a* (NCBI ref. # 117508) in B3 with B3 Alexa Fluor 647.

Whole embryo immunohistochemistry was performed as described in Subramanian et al., 2018. Primary antibodies used: rat monoclonal anti-mCherry (Molecular Probes − 1:500 dilution, M11217, RRID: AB_2536611), chicken anti-GFP (Abcam – 1:1000 dilution, ab13970, RRID:AB_300798), mouse anti-myosin heavy chain (MHC) (Developmental Hybridoma - 1:250, A1025, RRID:AB_528356). Secondary antibodies used: Alexa Fluor 594 conjugated donkey anti-rat IgG (Jackson ImmunoResearch – 1:1000 dilution, 712-586-153, RRID:AB_2340691), Alexa Fluor 488 conjugated donkey anti-chicken IgY (Jackson Immunoresearch, 1:1000 dilution, 703-486-155, RRID:AB_2736851), Alexa Fluor 647 conjugated donkey anti-mouse IgG (Jackson Immunoresearch, 1:1000 dilution, 715-606-151, RRID:AB_2340866).

### Embryo dissociation and FACS sorting

For comparing 36-48 hpf bulk RNA-seq, transgenic *Tg*(*scxa:mCherry*) zebrafish embryos were dissociated using collagenase IV (Gibco, 17104019) at a concentration of 6.25 mg/ml without trypsin addition at a temperature of 28°C for roughly 40 minutes, homogenizing every 5 min using a P1000 pipette as described in (Barske et al., 2016). Cells were then filtered through a 40 μm filter (Pluriselect-usa, 43-10040-50). Dissociated cell suspensions were sorted on a Bio-Rad FACS Aria II cell sorter. mCherry-positive cells were gated and sorted for those expressing at high levels.

For aBTX injected 48 hpf bulk RNA-seq alone transgenic *Tg*(*scxa:mCherry*) embryos, BTX-injected or un-injected siblings, were dissociated using Subtilisin A cold-active protease in a stock solution consisting of: 5 ul of 1M CaCl2, 100 ul of protease stock solution (100mg of Bacillus licheniformis protease (Sigma P5380) solubilized in 1 ml of Ca and Mg free PBS), 889ul of PBS, 1 ul of 0.5M EDTA (Sigma E5134) and 5ul of DNAse I (Roche 10104159001) stock (25U/ul in PBS, stored at -80C) adapted from (O’Flanagan et al., 2019). Embryos were triturated once every 2 min for 15 seconds using a wide bore 1 ml pipette. Every 15 min, tissue solution as checked under dissecting scope to verify dissociation. Full dissociation took ∼30 min per samples, and samples were subsequently run through a 40 μm filter to separate dissociated cells from clumps of aggregate undissociated tissue/ECM and washed with 10 ml of PBS/BSA (0.01% BSA in PBS, made fresh on the day of dissociation) and transferred to a 15 ml conical tube. Cells were centrifuged at 600g for 5 min at 4°C, supernatant is discarded, and cells are resuspended in 1 ml of ice-cold PBS/BSA before being placed on ice. **(Manuscript to be submitted).** High expressing mCherry+ cells were gated and sorted on a Bio-Rad FACS Aria II cell sorter.

### Bulk RNA-seq library preparation and sequencing

For comparing 36-48 hpf bulk RNA-seq samples an RNEasy Micro Kit (Qiagen, 74004) was used for RNA extraction of cell lysates from FAC-sorted cells. RNA quality was checked at the UC Irvine Genomics High Throughput Facility (GHTF) using a Bioanalyzer 2100 Instrument (Agilent). The Smart-seq2 protocol was utilized for cDNA library construction (Picelli et al., 2014). Libraries were sequenced at the GHTF using a HiSeq 4000 sequencer (Illumina) at a read depth of ∼35M reads per replicate. From 11 total biological replicates (7 for 36 hpf, 4 for 48 hpf) we obtained approximately 10,000 cells per sample replicate.

For 48 hpf bulk RNA-seq library preparations from aBTX-injected and un-injected siblings were performed by the UCI GHTF. Libraries were sequenced at GHTF on a NextSeq 550 sequencer (Illumina) at a read depth of ∼35M reads per replicate

### Bulk RNA-seq data analysis

Reads were mapped to zebrafish genome version GRCz10 and quantified using STAR v2.5.2a (Dobin et al., 2013) and RSEM v1.2.31 (B. Li & Dewey, 2011). Differential gene expression analysis and PCA were performed using R package DESeq2 v1.30.1 (Love et al., 2014). Pairwise comparisons were performed between 36 hpf and 48 hpf sorted tenocytes, and a Benjamini-Hochberg FDR adjusted p-value < 0.05 was used as a threshold for considering significant differences in gene expression levels. PCA was performed on normalized count data which underwent variance-stabilization-transformation using DESeq2. Heatmaps were generated using ClustVis (Metsalu & Vilo, 2015). GO term enrichment analysis was performed using the ClusterProfiler R package (T. Wu et al., 2021) and ShinyGO (Ge et al., 2020).

### **α**BTX injections

αBTX mRNA was synthesized from the *Pmtb-t7-alpha-bungarotoxin* vector (Megason lab, Addgene, 69542) as described in (Swinburne et al., 2015) and injected into embryos at the 1-cell stage at a volume of 500 picoliters per embryo. αBTX mRNA was injected at a concentration of 90 ng/μl (45 pg)to paralyze embryos that were collected at 48 hpf and 150 ng/μl (90 pg) to paralyze embryos that were collected at 72 hpf.

### RT-qPCR

Wild type, *cacnb1^-/-^*, αBtx-paralyzed, twitching, and recovered embryos were collected at respective timepoints, homogenized in Trizol with prefilled tube kits using high impact zirconium beads (Benchmark Scientific, D1032-10) using a BeadBug 3 Microtube Homogenizer D1030 (Benchmark Scientific), and RNA was extracted as described earlier. cDNA was prepared according to the standard oligo-dT primer protocol using the ProtoScript II First Strand cDNA Synthesis Kit (NEB E6560). cDNA was diluted 1:25 in water and used as template for RT-qPCR using the Luna Universal qPCR master mix (NEB M3003S). Primers used are listed in **Supplementary File 7.** Primer efficiencies were calculated with the formula PCR-efficiency = 10^(−1/slope)^ from a linear regression of Cp/ln(DNA) using a serial dilution of each primer with 72 hpf embryo cDNA as described in (Pfaffl, 2001). PCR reactions were performed on a LightCycler 480 II Real Time PCR Instrument (Roche) and analyzed using LightCycler 480 Software. Each RT-qPCR experiment was repeated in triplicate for each biological replicate, and at least two biological replicates were used for each analysis. P-values were calculated using a two-tailed Student’s T-test with α = 0.05 in Microsoft Excel. Bar charts in **Fig. 3** present mean +/- standard error. Venn diagram was created using the VennDiagram v1.7.3 R package with the gene list overlap tested with the Fishers exact test from the GeneOverlap v1.26.0 R package (Li Shen, 2017).

### Imaging and *is*HCR quantification

Whole embryos imaged for ISH were mounted on slides in 80% glycerol and imaged using a Zeiss Axioplan 2 compound microscope utilizing an AxioCam 305 Color Micropublisher 5.0 RTV camera with Zeiss Zen 3.1 (blue edition) software. Embryos imaged for *is*HCR were embedded in 1% low melting point agarose/5x SSC and imaged on a Leica SP8 confocal microscope using the PL APO CS2 40X/1.10 W objective. Whole embryos imaged for *is*HCR were mounted in slide dishes in 1% low melt agarose with either 5xSSC (if only *is*HCR was performed) or 1xPBT (if *is*HCR with immunofluorescence was performed) and imaged using a Leica SP8 confocal microscope with LASX software. *is*HCR voxel colocalizations in **Fig. 2** and **Fig. 2—supplemental figure 1** were performed using the “Coloc” function in Imaris 10.0.1 as described in (Subramanian et al., 2023). Voxel colocalization only shows overlap of fluorescent channels within a particular voxel which may not, in some instances, fully reflect actual colocalization of fluorescence within a particular cell due to the punctate nature of *is*HCR fluorescence. *is*HCR single cell quantification was performed in Imaris 10.0.1 using DAPI as a nuclear marker, as described in (Subramanian et al. 2023). Embryo imaging for a single experiment was performed with identical parameters across conditions. Briefly, an ROI of the DAPI-stained nucleus from each 3D stack was traced through individual z-slices and mean voxel-intensity (AU) was measured. *matn1/scxa* co-expressing cells measured were located at the ima enthesis on meckels cartilage and sht enthesis at the anterior edge of the ch cartilage. *klf2a/scxa* and *mxra5b/scxa* co-expressing cells measured were located at the ima enthesis and sht enthesis, mhj MTJ and sht MTJ. *klf2a/scxa* co-expression was measured primarily in tenocytes near the horizontal myoseptum (HMS) whereas *mxra5b/scxa* co-expression was measured primarily from tenocytes in the ventral half of the vertical myoseptum (VMS). Experimental conditions pertaining to each embryo image were saved separately, measurements were performed, and conditions were matched to each image. All p-values were calculated using a linear mixed effects model with individual embryos set as the random variable, and cells set as the fixed variable using the lme4 and lmetest packages. Tukey-Kramer post-hoc tests for pairwise analyses where then performed (ns = not significant, * p < 0.05, ** p < 0.01, *** p < 0.001).

### Multiplex CRISPR-Cas9 genome editing of *matn1*, *klf2a*, and *mxra5b*

*matn1, klf2a,* and *mxra5b* multiplex gRNA injections were performed using the methodology described in (T. Wu et al., 2021) using gRNA primer sequences were obtained from the primer database provided. Briefly, PCR was performed with four primers (per gene) targeting coding regions with T7 and spacer sequences for template gRNA synthesis. Transcription was performed with the T7 Megashortscript kit (Invitrogen AM1354). A 500 ng/ul solution of all 4 gRNAs were incubated at 37°C and injected into 1-cell stage embryos at a 500 picoliter volume per embryo.

## Acknowledgements

We would like to acknowledge Dr. Daniel Dranow for reviewing the manuscript and assistance provided for experimental design. This study was made possible in part through access to the Optical Biology Core Facility of the Developmental Biology Center, a shared resource supported by the Cancer Center Support Grant (CA-62203)

## Author contributions

Conceptualization: P.K.N., A.S., T.F.S.; Methodology: P.K.N., A.S.; Validation: P.K.N., A.S.; Formal analysis: P.K.N., A.S.; Investigation: P.K.N., A.S.; Resources: T.F.S.; Writing - original draft: P.K.N..; Writing - review & editing: P.K.N., A.S., T.F.S.; Visualization: P.K.N.; Supervision: T.F.S.; Project administration: T.F.S.; Funding acquisition: T.F.S.

---

**Figure 1—figure supplement 1:**
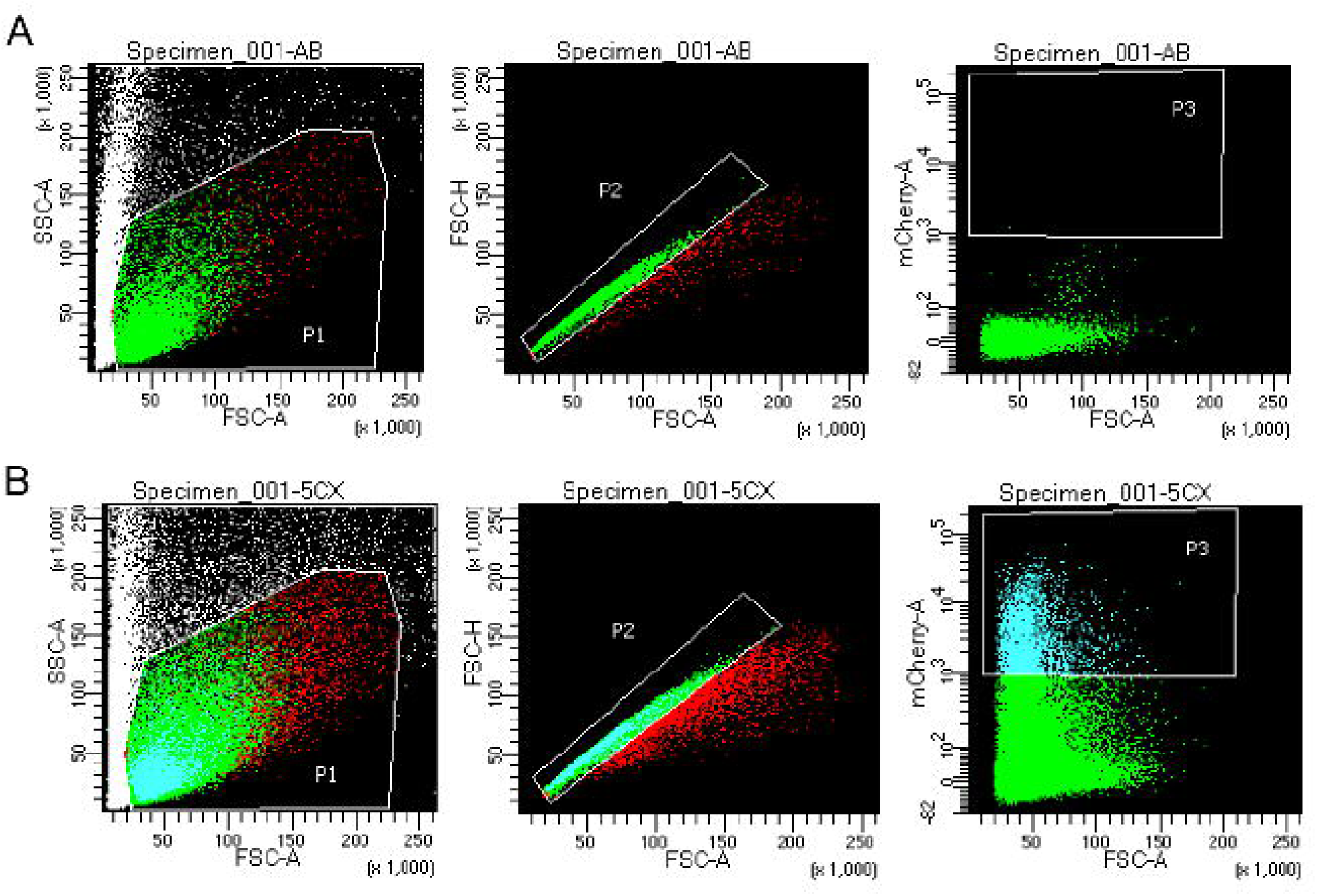
FACS Gating thresholds for mCherry+ cells. (Left to right for each) Forward Scatter A (FSC-A) versus Side Scatter A (SSC-A) shows P1 threshold. FSC-A versus Forward Scatter H (FSC-H) shows P2 threshold to select for single cells. FSC-A versus mCherry-A shows P3 fluorescence gating for mCherry+ cells. **A)** P3 selection gating allowed selection of cells with strong mCherry signal based on Negative control AB (WT) sample vs mCherry expressing tenocytes (48 hpf) FACS gating. Established P3 gating selected for mCherry positive cells in all 36 hpf and 48 hpf *Tg(scxa:mCherry)* samples. **B)** Thresholds used in 36 hpf and 48 hpf *Tg(scxa:mCherry)* samples for FACS prior to bulk RNA-seq.

**Figure 2—figure supplement 1:**
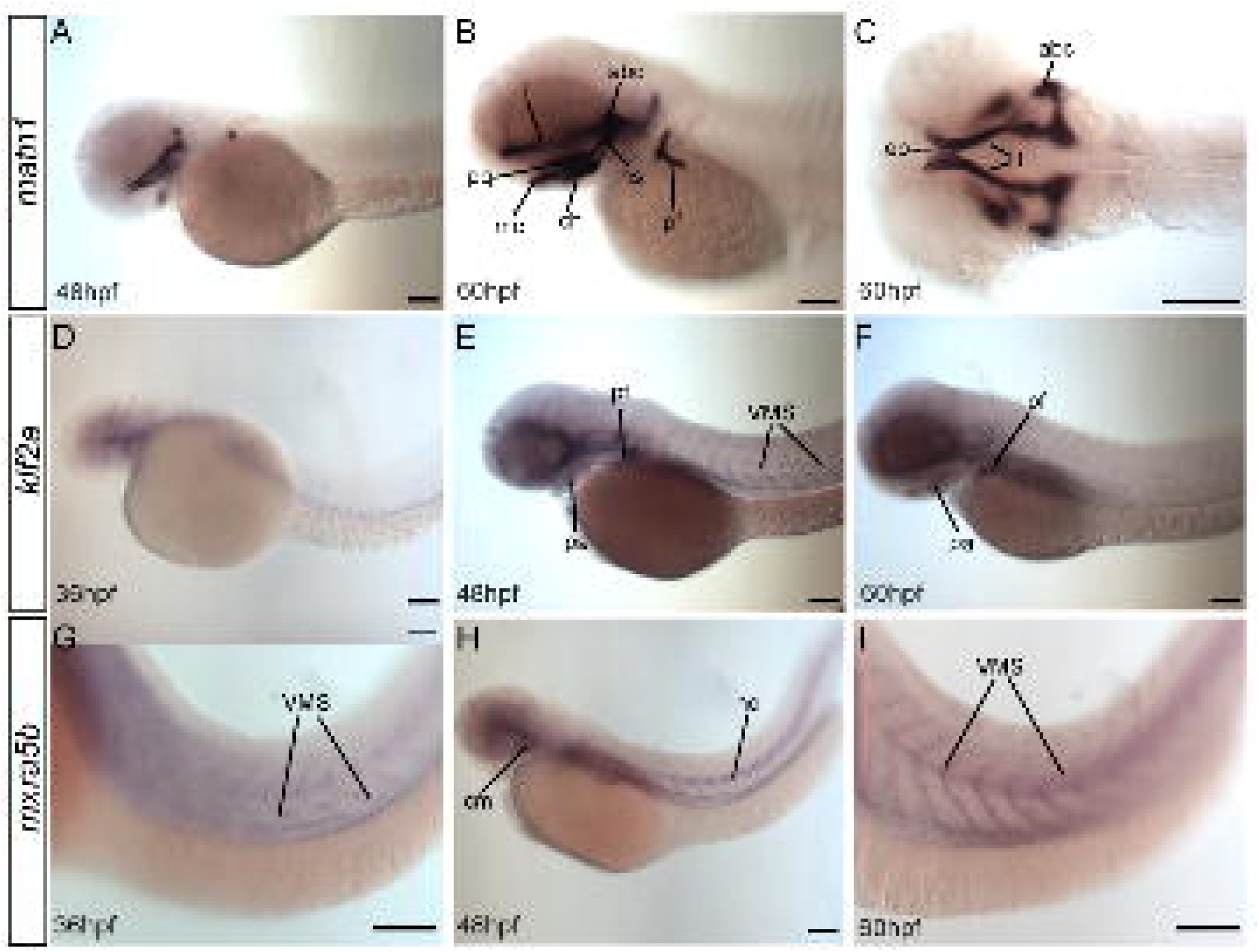
*matn1, klf2a, mxra5b* are expressed in musculoskeletal tissues of developing embryos. Lateral **(A,B, D-I)** and ventral **(C)** views of embryos showing expression of *matn1* **(A-C)**, *klf2a* **(D-F)** and *mxra5b* **(G-H) (A-C)** 48 hpf embryos show *matn1* expression in cartilage progenitors at 48 hpf and in differentiated cartilages (and associated tenocytes) at 60 hpf **(B,C)**. **(D-F)** Lateral views of 36 hpf **(D)**, 48 hpf **(E)** and 60 hpf **(F)** embryos show *klf2a* expression in pharyngeal mesenchyme **(D)**, skeletal progenitors and in tenocytes along vertical myosepta (VMS) **(E,F)**. **(G-I)** Lateral views of 36 hpf **(G)**, 48 hpf **(H)** and 60 hpf **(I)** embryos show *mxra5b* expression in tenocytes along horizontal myosepta (HMS) along the notochord and VMS. Scale bars = 100μm. Abbreviations: abc = anterior basicranial commissure, ch = ceratohyal cartilage, ep = ethmoid plate, hs = hyosymplectic cartilage, mc = meckel’s cartilage, nc = notochord, pf = pectoral fin, pq = palatoquadrate cartilage, sb = somite boundaries, t = trabeculae cartilage.

**Figure 2—figure supplement 2:**
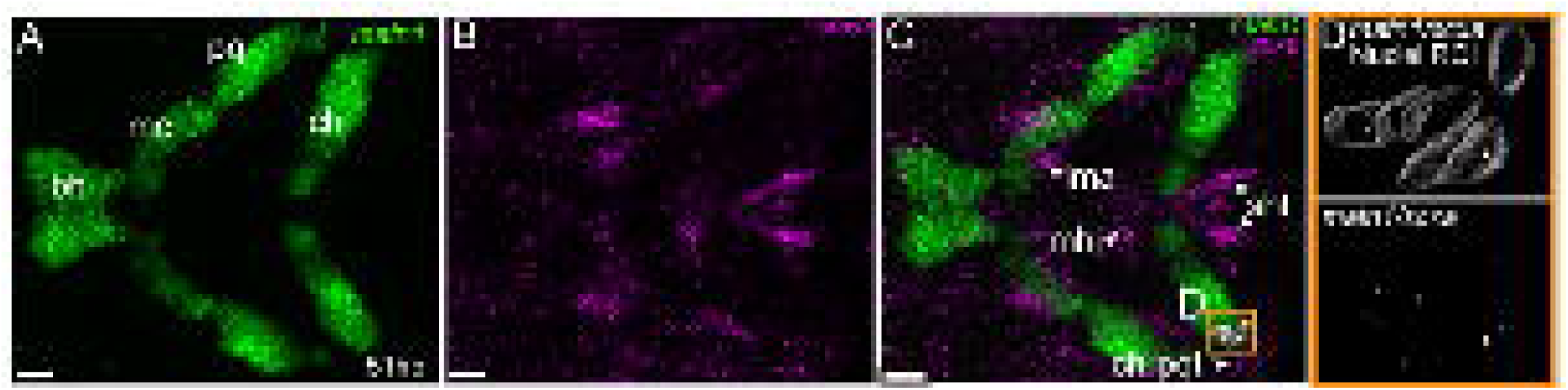
*matn1* is expressed in differentiating cranial tenocytes. Ventral view of the developing mandibular arch in a 51 hpf embryo showing *is*HCR of *matn1* **(A,C,D)** and *scxa* **(B-D)**. **D)** magnified view of yellow ROI **(C)** shows outline of tenocyte nuclear 3D-volume with white puncta representing voxel colocalizations of *matn1* and *scxa* as depicted by colocalization using Imaris (see methods). bh- basihyal cartilage, ch = ceratohyal cartilage, mc = meckel’s cartilage, pq = palatoquadrate cartilage, ch-pqt – ceratohyal-palatoquadrate tendon, ima = intermandibularis anterior tendon, mhj = mandibulohyoid junction tendon, sht = sternohyoideus tendon. Scale bars = 20 microns.

**Figure 3—figure supplement 1:**
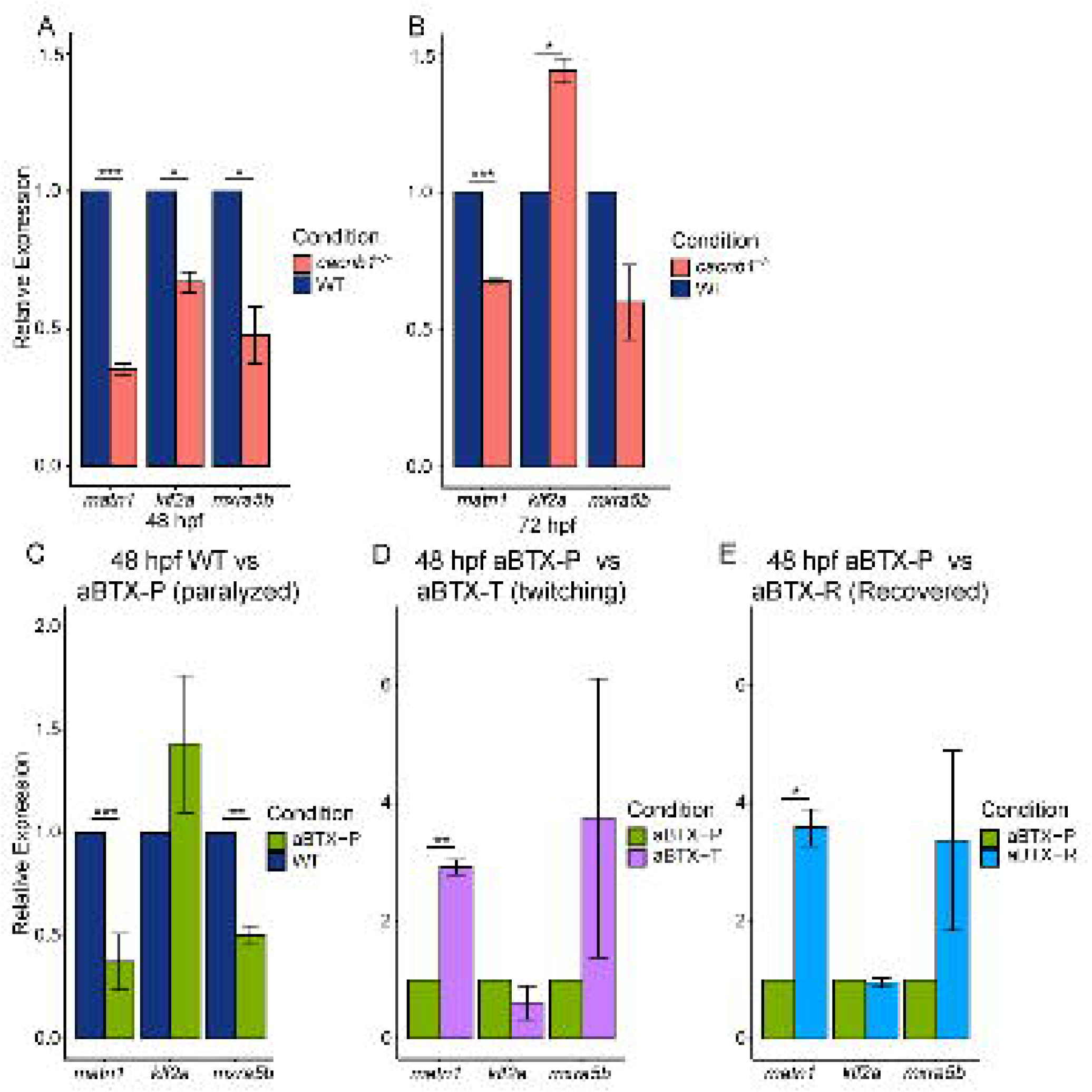
Paralysis regulates gene expression of *matn1, klf2a,* and *mxra5b* in developing embryos. **(A,B)** Bar plots showing global changes in relative expression from RT-qPCR of *matn1, klf2a,* and *mxra5b* genes between WT and *cacnb1^-/-^* mutant embryos at 48 hpf **(A)** and 72 hpf **(B)**. **(C-E)** Bar plots show global changes in relative expression of *matn1*, *klf2a*, and *mxra5b* between 48 hpf un-injected WT controls (blue) and α*BTX*-injected paralyzed (green) embryos **(C)**, α*BTX*- injected paralyzed (green) and αBTX-injected “Twitching” (partially recovered, magenta) embryos **(D)**, and between α*BTX*-injected paralyzed (green) and αBtx-injected, “Recovered” (cyan) embryos **(E)** at 48 hpf (right). ns = not significant, * p < 0.05, ** p < 0.01, *** p < 0.001

**Figure 4—figure supplement 1:**
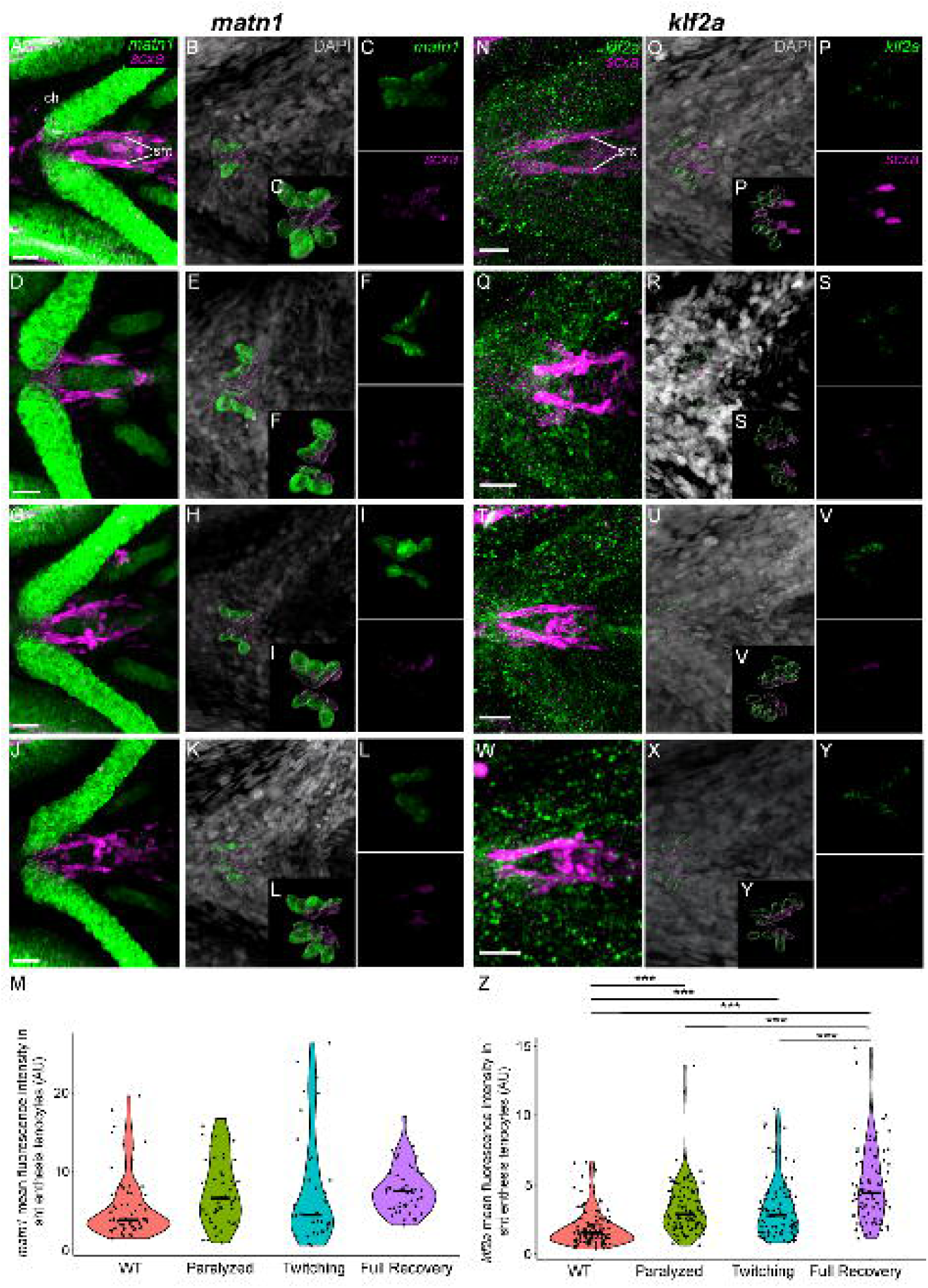
Mechanical force regulates expression of *matn1* and *klf2a* in sh-ch (sht) enthesis tenocytes. Ventral views of ceratohyal (ch) cartilage and associated tenocytes showing *is*HCR of *matn1* (green) **(A-L)** and *klf2a* (green) **(N-Y)** and anti-mCherry immunofluorescence (magenta) marking the tenocytes in *Tg(scxa:mCherry)* embryos at 72 hpf in WT uninjected (WT) **(A-C, N-P)**, *αBTX*- inj paralyzed **(D-F, Q-S)**, partially recovered *αBTX*-inj (Twitching) **(G-I, T-V)**, and completely recovered *αBTX*-inj (Full Recovery) **(J-L, W-Y)** conditions at sht enthesis. **(B, E, H, K, O, R, U, X)** Grayscale images showing nuclei stained with DAPI with ROIs showing isolated 3D-volumes of chondrocytes (green) and sternohyoideus-ceratohyal (sh-ch) sht enthesis tenocytes (magenta) based on DAPI signal. **(C, F, I, L, P, S, V, Y)** Insets showing magnified views of the 3D-volumes of tenocytes associated with sh-ch enthesis depicting expression of *matn1* and stained for mCherry. **(M)** Violin plot showing changes in mean fluorescence intensity of *matn1* in sh-ch enthesis tenocyte nuclei between WT (n=8), Paralyzed (n=8), Twitching (n=6) and full Recovery (n=7) with ∼8 nuclei measured per embryo. **(Z)** Violin plot showing changes in mean fluorescence intensity of *klf2a* in sh-ch (sht) enthesis tenocyte nuclei between WT (n=15), paralyzed (n=16), twitching (n=14) and full Recovery (n=11) with ∼8 nuclei measured per embryo. p-values calculated with linear mixed effects model with Tukey post-hoc test. * p < 0.05, ** p < 0.01, *** p < 0.001). Scale bars = 20 microns. Figure 4—figure supplement 1 source data 2 Measurements of *matn1* and *klf2a* HCR signal intensity in sht enthesis tenocytes [Figure 4— figure supplement 1 Source Data.xlsx]

**Figure 5—figure supplement 1:**
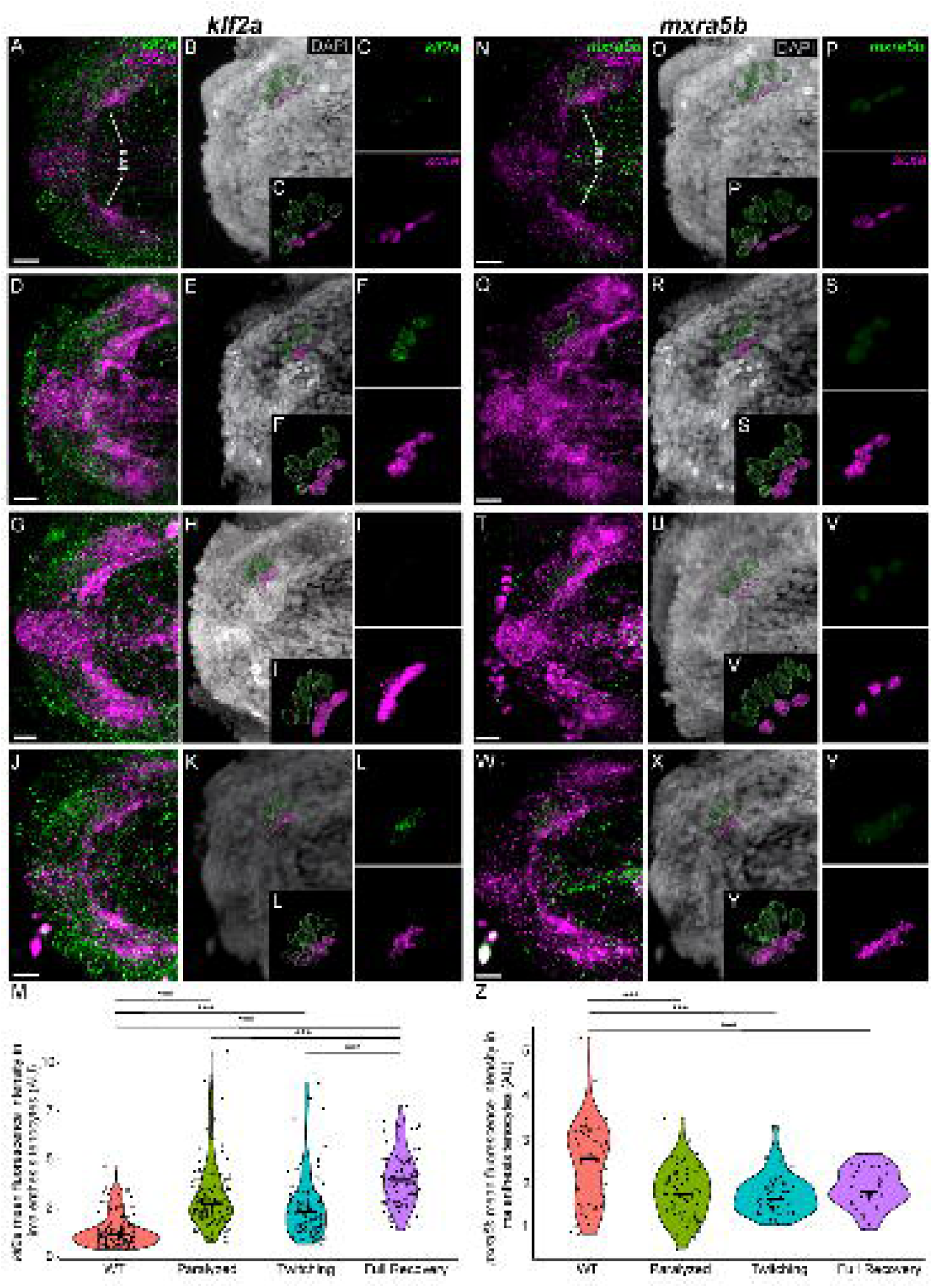
Mechanical force differentially regulates expression of *mxra5b* and *klf2a* in ima enthesis tenocytes. Ventral views of Meckels cartilage and associated tenocytes showing *is*HCR of *klf2a* (green) **(A-L)** and *mxra5b* (green) **(N-Y)** and anti-mCherry immunofluorescence (magenta) marking the tenocytes in *Tg(scxa:mCherry)* embryos at 72 hpf in WT uninjected (WT) **(A-C, N-P)**, *αBTX*-inj paralyzed **(D-F, Q-S)**, partially recovered *αBTX*-inj (Twitching) **(G-I, T-V)**, and completely recovered *αBTX*-inj (Full Recovery) **(J-L, W-Y)** conditions at ima enthesis. **(B, E, H, K, O, R, U, X)** Grayscale images showing nuclei stained with DAPI with ROIs showing isolated 3D-volumes of chondrocytes (green) and ima enthesis tenocytes (magenta) based on DAPI signal. **(C, F, I, L, P, S, V, Y)** Insets showing magnified views of the 3D-volumes of tenocytes associated with ima enthesis depicting expression of *mxra5b* and *klf2a* and stained for mCherry. **(M)** Violin plot showing changes in mean fluorescence intensity of *klf2a* in ima enthesis tenocyte nuclei between WT (n=15), Paralyzed (n=16), Twitching (n=14) and Full Recovery (n=11) with ∼8 nuclei measured per embryo. **(Z)** Violin plot showing changes in mean fluorescence intensity of *mxra5b* in ima enthesis tenocyte nuclei between WT (n=7), Paralyzed (n=8), Twitching (n=8) and Full Recovery (n=4) with ∼8 nuclei measured per embryo. p-values calculated with linear mixed effects model with Tukey post-hoc test. * p < 0.05, ** p < 0.01, *** p < 0.001). Scale bars = 20 microns. Figure 5—figure supplement 1 source data 4 Measurements of *kfl2a* and *mxra5b* HCR signal intensity in ima enthesis tenocytes [Figure 5— figure supplement 1 Source Data.xlsx]

**Figure 5—figure supplement 2:**
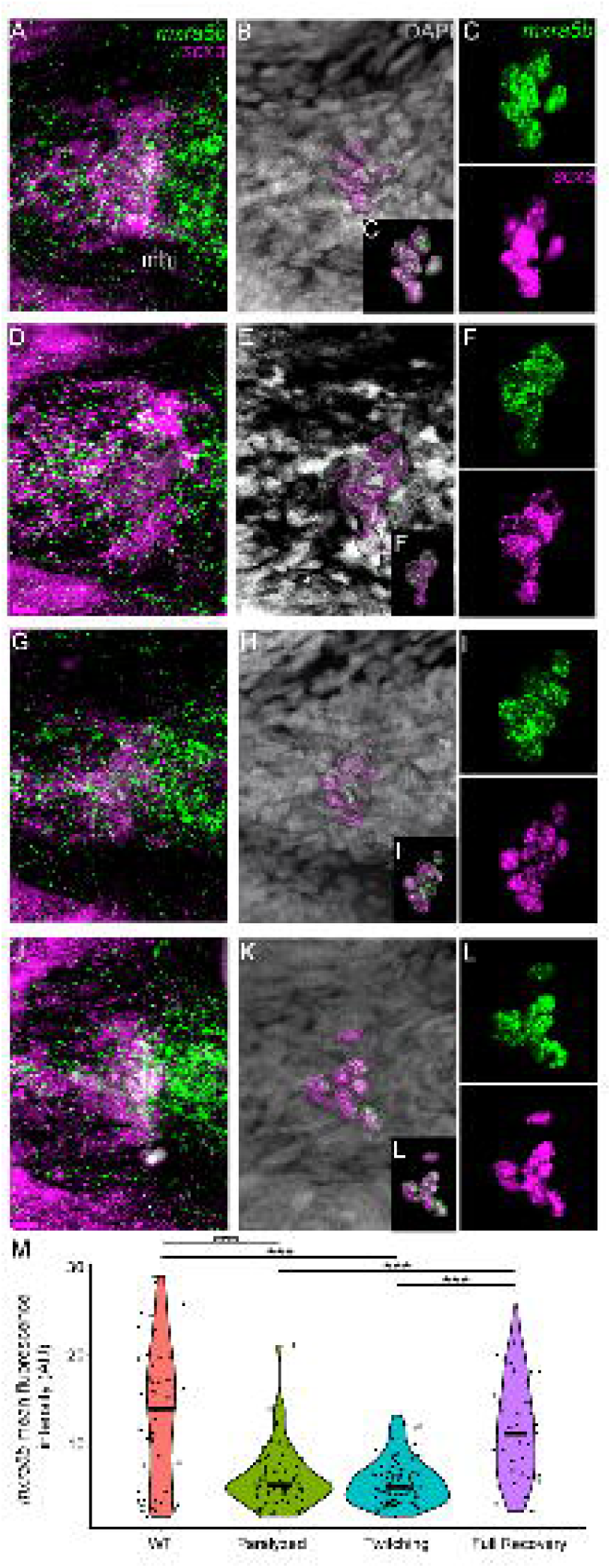
Mechanical force regulates expression of *mxra5b* in mhj myotendinous tenocytes. Ventral views of mandibulohyoid(mhj) myotendinous junction (mtj) associated tenocytes showing *is*HCR of *mxra5b* (green) and anti-mCherry immunofluorescence (magenta) marking the tenocytes in *Tg(scxa:mCherry)* embryos at 72 hpf in WT uninjected (WT) **(A-C)**, *αBTX*-inj paralyzed **(D-F)**, partially recovered *αBTX*-inj (Twitching) **(G-I)**, and completely recovered *αBTX*- inj (Full Recovery) **(J-L)** conditions. **(B,E,H,K)** Grayscale images showing nuclei stained with DAPI with ROIs showing isolated 3D-volumes of mhj tenocytes (magenta) based on DAPI signal. **(C,F,I,L)** Insets showing magnified views of the 3D-volumes of tenocytes associated with mhj-mtj depicting expression of *mxra5b* and stained for mCherry. **(M)** Violin plot showing changes in mean fluorescence intensity of *mxra5b* in mhj-mtj tenocyte nuclei between WT (n=7), Paralyzed (n=8), Twitching (n=8) and Full Recovery (n=4) with ∼10 nuclei measured per embryo. p-value calculated with linear mixed effects model with Tukey post-hoc test. * p < 0.05, ** p < 0.01, *** p < 0.001). Scale bars = 20 microns. Figure 5—figure supplement 2 source data 5 Measurements of *mxra5b* HCR signal intensity in mhj mtj tenocytes [Figure 5—figure supplement 2 Source Data.xlsx]

**Figure 5—figure supplement 3:**
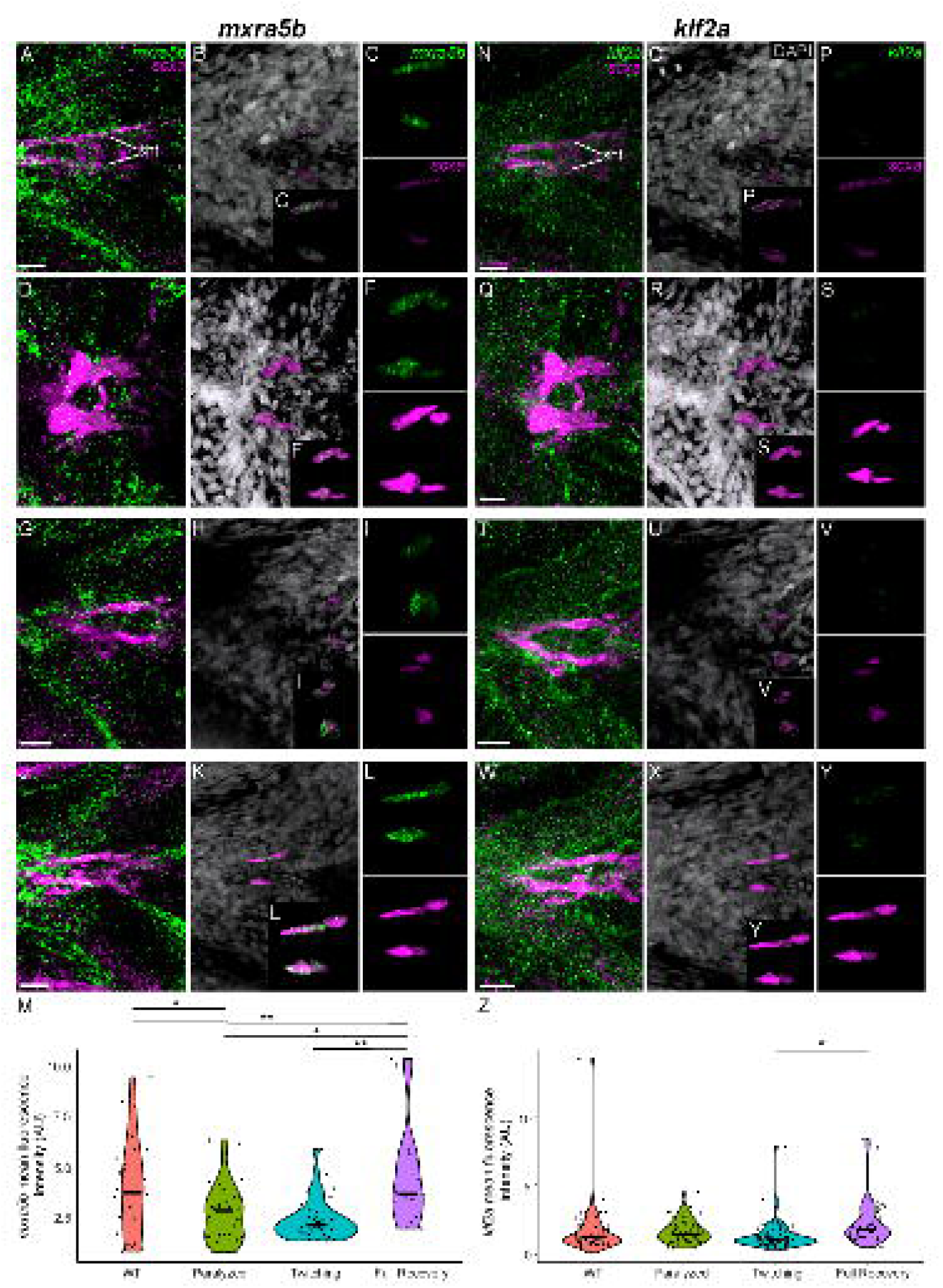
Mechanical force differentially regulates expression of *mxra5b* and *klf2a* in sht myotendinous junction tenocytes. Ventral views of sternohyoideus tendon and associated tenocytes showing *is*HCR of *mxra5b* (green) **(A-L)** and *klf2a* (green) **(N-Y)** and anti-mCherry immunofluorescence (magenta) marking the tenocytes in *Tg(scxa:mCherry)* embryos at 72 hpf in WT uninjected (WT) **(A-C, N-P)**, *αBTX*- inj paralyzed **(D-F, Q-S)**, partially recovered *αBTX*-inj (Twitching) **(G-I, T-V)**, and completely recovered *αBTX*-inj (Full Recovery) **(J-L, W-Y)** conditions at sht MTJ. **(B, E, H, K, O, R, U, X)** Grayscale images showing nuclei stained with DAPI with ROIs showing isolated 3D-volumes of ima enthesis tenocytes (magenta) based on DAPI signal. **(C, F, I, L, P, S, V, Y)** Insets showing magnified views of the 3D-volumes of tenocytes associated with ima enthesis depicting expression of *mxra5b* and *klf2a* and stained for mCherry. **(M)** Violin plot showing changes in mean fluorescence intensity of *mxra5b* in sht MTJ tenocyte nuclei between WT (n=7), Paralyzed (n=8), Twitching (n=8) and Full Recovery (n=4) with ∼4 nuclei measured per embryo. **(Z)** Violin plot showing changes in mean fluorescence intensity of *klf2a* in ima enthesis tenocyte nuclei between WT (n=15), Paralyzed (n=16), Twitching (n=14) and Full Recovery (n=11) with ∼4 nuclei measured per embryo. p-values calculated with linear mixed effects model with Tukey post-hoc test. * p < 0.05, ** p < 0.01, *** p < 0.001). Scale bars = 20 microns. Figure 5—figure supplement 3 source data 6 Measurements of *mxra5b* and *klf2a* HCR signal intensity in sht mtj tenocytes [Figure 5—figure supplement 3 Source Data.xlsx]

